# Kinesin-5/Cut7 moves bidirectionally on fission-yeast spindles with activity that increases in anaphase

**DOI:** 10.1101/2022.08.01.502387

**Authors:** Zachary R. Gergely, Saad Ansari, Michele H. Jones, J. Richard McIntosh, Meredith D. Betterton

## Abstract

Kinesin-5 motors are essential to separate mitotic spindle poles and assemble a bipolar spindle in many organisms. These tetrameric motors crosslink and slide apart antiparallel microtubules via microtubule plus-end-directed motility. However, kinesin-5s typically accumulate more at spindle poles than in the center of the spindle where antiparallel microtubule overlaps are most numerous. While the relevance of this localization has remained unclear, increasing evidence suggests that it occurs due to bidirectional motility or trafficking of kinesin-5 motors. The kinesin-5 motor Cut7p from fission yeast has been shown to move bidirectionally in reconstituted systems. However, bidirectional movement has not been shown in cells and the funtion of the minus-end-directed movement remains unclear. Here, we characterized the motility of kinesin-5/Cut7 on bipolar and monopolar spindles in fission yeast and observed movement both toward plus and minus ends of microtubules. Notably, we found that the activity of the motor increases at the onset on anaphase B. Perturbations to microtubule dynamics did not significantly change the observed Cut7p movement, while Cut7p mutation reduced or abolished observable movement. These results suggest that the directed movement of Cut7p was due to the motility of the motor itself. Mutations of Cut7p that decreased plus-end-directed motility enhanced its spindle-pole localization. In contrast, abolishing Cut7 motility or replacing it with plus-end-directed human Eg5 eliminates the pole localization. Our results suggest a new hypothesis for the function of minus-end-directed motility and spindle-pole localization of kinesin-5s in spindle assembly.

## INTRODUCTION

While kinesin-5 motors have long been known to play an important role in bipolar spindle assembly and mitotic chromosome segregation, the links between kinesin-5 motility, localization, and force generation remain incompletely understood. Kinesin-5 motors are essential for mitotic spindle formation in many organisms because they separate spindle poles to build a bipolar spindle (Enos and Morris (1990); Hagan and Yanagida (1990); Hoyt et al. (1992); Sawin et al. (1992); Blangy et al. (1995), Fig. 1). Kinesin-5s are homo-tetramers, with two dimeric motors linked antiparallel by a central minifilament (Cole et al. (1994); Kashina et al. (1996); Gordon and Roof (1999); Acar et al. (2013); Scholey et al. (2014); Singh et al. (2018), Fig. 1A,B). Thus kinesin-5 can crosslink antiparallel microtubules (MTs) and slide them apart, both in cells and in reconstituted systems (Sharp et al., 1999; Kapitein et al., 2005; Hildebrandt et al., 2006; Tao et al., 2006; van den Wildenberg et al., 2008; Shimamoto et al., 2015; Bodrug et al., 2020). This activity contributes both to spindle pole separation as the bipolar spindle forms and to spindle elongation in anaphase B (Goshima and Scholey, 2010; Mann and Wadsworth,2018; Scholey et al., 2016). Antiparallel sliding is therefore crucial to kinesin-5 function in mitosis, and depends on the motor’s motility toward MT plus ends (Fig. 1C). Consistent with this view, kinesin-5 depletion or inhibition leads to monopolar spindles (Hagan and Yanagida, 1990; Sawin et al., 1992; Mayer et al., 1999).

**Fig. 1.**
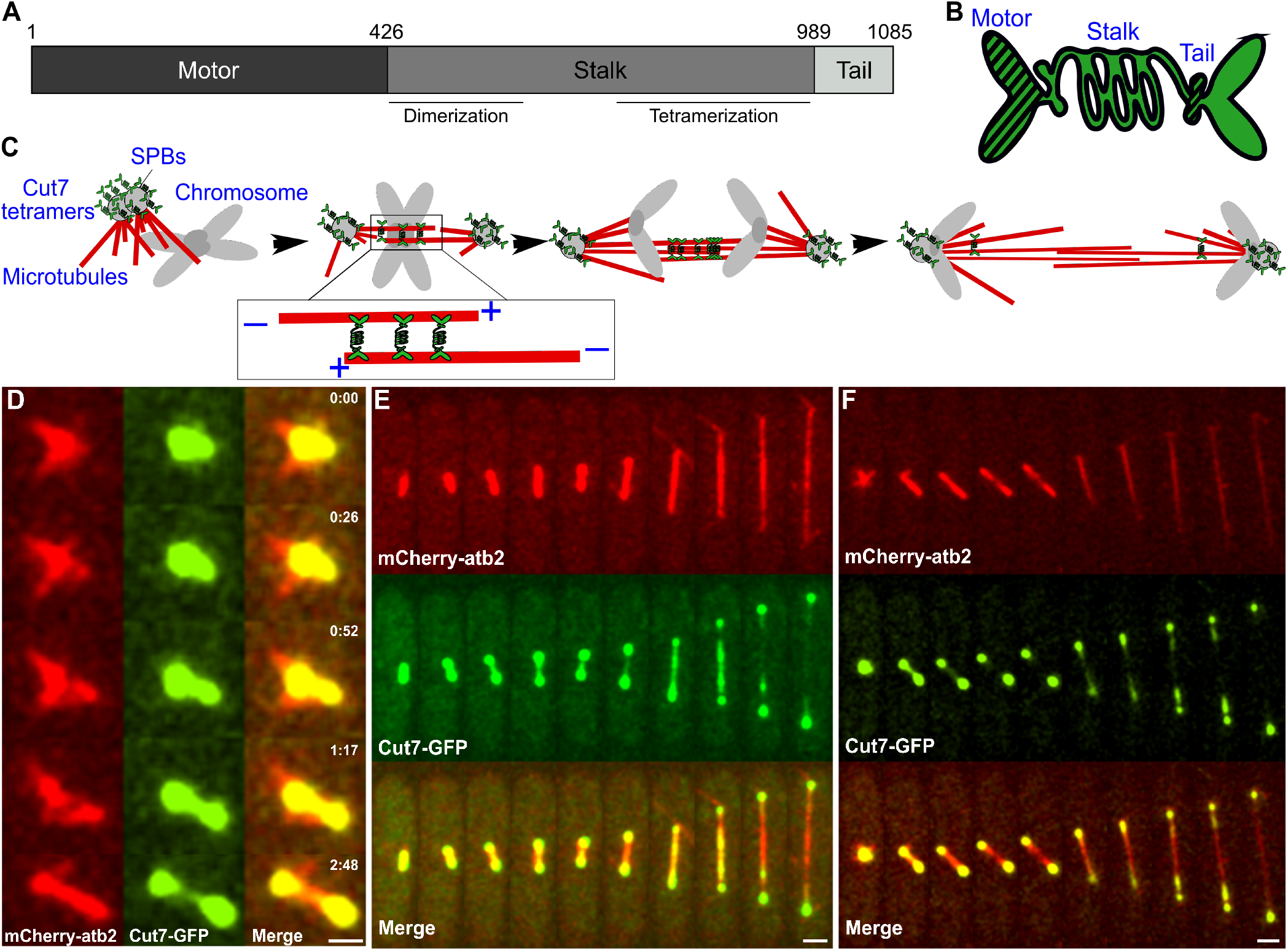
Overview of Cut7p structure and localization (A) The major domains of Cut7p delimited by amino acid locations. (B) Cartoon of a Cut7p tetramer, showing its 2-fold symmetry. (C) Graphic showing localization of Cut7p tetramers on a mitotic spindle in S. pombe. Inset shows Cut7p crosslinking antiparallel MTs. (D) Cut7-GFP spindle localization in a temperature-sensitive cut11-7 strain at permissive temperature. As the monopolar spindle becomes bipolar, a subset of Cut7-GFP (green) moves from the MTs (red) near the left SPB to the MTs near the right SPB. Scale bar: 2 *μ*m. (E-F) Example of cut11+ (E) and cut11-7 (F) cells showing Cut7-GFP (green) localization on MTs (red) throughout mitosis. All images taken at 25°C. Scale bars: 1 *μ*m.

However, more recent results have called into question whether the elegantly simple antiparallel-sliding model can fully explain kinesin-5 function. Antiparallel sliding requires antiparallel MTs, which are most abundant near the center of the spindle, where microtubules from both poles interdigitate (McIntosh and Landis, 1971; Ding et al., 1993). However, kinesin-5s localize more strongly near spindle poles in many organisms (Fig. 1), including *Schizosaccharyomyces pombe* (Hagan and Yanagida, 1992), *Saccharomyces cerevisiae* (Shapira et al., 2017), *Xenopus laevis* (Sawin et al., 1992; Cahu et al., 2008), and mammalian (Blangy et al., 1995; Gable et al.,2012) cells. We have limited understanding of how pole localization occurs for a plus end-directed motor, but there are three broad ideas for the mechanism: trafficking by a minus-end-directed motor that carries kinesin-5 as cargo, intrinsic bidirectional motility of kinesin-5 that moves motors toward spindle poles, and direct pole binding.

Minus-end-directed trafficking occurs for vertebrate kinesin-5/Eg5, which is plus-end directed but can be transported toward MT minus ends by dynein (Uteng et al., 2008; Gable et al., 2012) The trafficking is regulated by TPX2 (Ma et al., 2010; 2011; Gable et al., 2012; Balchand et al., 2015). Importantly, chimeric kinesin-5s made from the Eg5 tail and the motor of other kinesins fail to assemble a bipolar spindle, even though these motors localize to spindle MTs (Cahu and Surrey, 2009). This evidence suggests that bidirectional trafficking of Eg5 by dynein may play a role in spindle assembly.

Both budding- and fission-yeast kinesin-5s have intrinsic bidirectional motility on MTs *in vitro* (Roostalu et al., 2011; Gerson-Gurwitz et al., 2011; Avunie-Masala et al., 2011; Thiede et al., 2012; Fridman et al., 2013; Edamatsu, 2014; 2016; Britto et al., 2016). For budding-yeast kinesin-5/Cin8, the average direction of motion changes with solution conditions and mutations (Gerson-Gurwitz et al., 2011; Roostalu et al., 2011; Shapira and Gheber, 2016; Shapira et al., 2017). Other work suggests that crowding of the MT lattice and motor clustering affect kinesin-5 speed and directionality (Britto et al., 2016; Shapira et al., 2017; Bodrug et al., 2020).

The kinesin-5 C-terminal tail contributes to spindle and spindle-pole localization. The tail contains conserved phosphorylation sites (Heck et al., 1993; Sawin and Mitchison, 1995; Blangy et al., 1995; Drummond and Hagan, 1998; Cahu et al., 2008; Rapley et al., 2008;Akera et al., 2015). Truncation or mutation of the tail can decrease or eliminate spindle (Sawin and Mitchison, 1995) or pole (Sharp et al., 1999; Olmsted et al., 2014) localization. Further, tail truncation in the metazoan kinesin-5s Eg5 and KLP61F significantly impairs microtubule crosslinking and sliding (Weinger et al., 2011; Bodrug et al., 2020), leading to greatly reduced sliding force generation (Bodrug et al., 2020). In budding yeast, deletion of the kinesin-5/Cin8 tail is lethal if the second kinesin-5/Kip1 is absent (Hildebrandt et al., 2006), consistent with idea that tail deletion impairs the ability of Cin8 to separate spindle poles. In fission yeast, kinesin-5/Cut7 pole localization occurs at least in part by direct binding of the tail to γ-tubulin at spindle poles (Olmsted et al., 2014). Recent work has suggested a structural mechanism by which tail-motor interactions could slow ATP hydrolysis (Bodrug et al., 2020). However, the mechanisms by which the C-terminal tail and its phosphorylation affect kinesin-5 motors are incompletely understood.

Previous computational modeling of *S. pombe* spindle assembly suggested that kinesin-5/Cut7 bidirectionality may localize it at spindle poles for proper spindle assembly (Blackwell et al., 2017; Edelmaier et al., 2020). Simulated bipolar spindles only assembled when kinesin-5 moved bidirectionally. The minus-end directed movement enhanced the motors’ spindle-pole localization in the model, positioning them to generate force and separate spindle poles (Blackwell et al., 2017; Edelmaier et al., 2020). Experimental work on budding-yeast kinesin-5 reached a similar conclusion (Shapira et al., 2017). However, direct experimental test of these model predictions in fission yeast has been limited because kinesin-5/Cut7 bidirectional motility has previously been observed only for the purified motor (Edamatsu, 2014; 2016; Britto et al., 2016). Furthermore, we currently do not understand why (or whether) bidirectional motility and pole localization are beneficial in mitosis, and whether bidirectional trafficking/motility of kinesin-5 may be altered at different times in mitosis.

Cut7p in the fission yeast *Schizosaccharomyces pombe* is a useful model for an in-depth study of this bidirectional kinesin on the spindle. This kinesin-5 is well-established as intrinsically bidirectional *in vitro* (Edamatsu, 2014; 2016; Britto et al., 2016). Cut7p is the sole kinesin-5 motor in this organism. It is essential in most genetic backgrounds (Hagan and Yanagida, 1990), but as in other cells, its loss can be rescued by deletion of this cell’s two kinesin-14s (Pidoux et al., 1996; Troxell et al., 2001; Rincon et al., 2017; Yukawa et al., 2018; Lamson et al., 2020; Yukawa et al., 2020). Moreover, Cut7p shares many properties with kinesin-5s more generally, including its basic domain structure and localization (Fig. 1). Study of directional motility on the spindle can take advantage of spindle-pole body (SPB) insertion defects in *cut11-ts* cells (West et al., 1998), which lead to monopolar spindles in cells containing only one SPB at restrictive temperature (West et al., 1998; Akera et al., 2015). Monopolar spindles with their minus ends tethered at the SPB can therefore be used to unambiguously identify motor movement toward MT plus- or minus-ends. Further, *S. pombe* monopolar spindles can enter anaphase with spindle MT elongation occurring at similar speeds as in bipolar spindles (Masuda et al., 1992; Krüger et al., 2021), facilitating study of changes in kinesin-5 motility before and after anaphase onset. Fission yeast therefore allow study of significant aspects of kinesin-5 function in a cell type that is suitable for detailed analysis.

## RESULTS

### Cut7p localizes to mitotic spindle microtubules and poles in *cut11*+ and *cut11-7* cells

To study kinesin-5/Cut7 localization and motility on fission-yeast spindles, we constructed strains containing Cut7-GFP and low-level mCherry-tubulin (Snaith et al. (2010); Blackwell et al. (2016), Fig. 1[D-F], Methods). *S. pombe cut7* fused to 1 or 3 GFP molecules at its C-terminal tail can replace the endogenous motor (Fu et al., 2009). The GFP protein is functional for spindle assembly with no observed growth or mitotic defects, and its localization is similar to the endogenous untagged motor (Hagan and Yanagida, 1992). This is consistent with observations of other GFP-tagged kinesin-5 motors (Avunie-Masala et al., 2011; Shapira et al., 2017). Consistent with previous studies (Hagan and Yanagida, 1992; Yukawa et al., 2015), we observed Cut7-GFP localized to the unseparated spindle poles early in mitosis, and also to the interpolar spindle as mitosis progressed (Fig. 1C). We observed similar behavior in *cut11*+ cells (Fig. 1E), and in temperature-sensitive *cut11-7* cells (West et al., 1998; Akera et al., 2015) grown at permissive temperature (Fig. 1F).

### Cut7p moves bidirectionally on bipolar spindles with activity that increases in anaphase B

To better assess Cut7p localization and motility during mitosis, we constructed kymographs of Cut7-GFP on bipolar spindles (Fig. 2). Bright Cut7-GFP signal was present near the spindle poles throughout mitosis (Fig. 2, S1), allowing the ends of the spindle to be easily identified in kymographs (Fig. 2A-H). As the spindle poles separated, bright clusters of Cut7-GFP appeared on the interpolar spindle. Diagonal streaks of Cut7-GFP signal were visible in the kymographs, which represent directional movement of one or more Cut7 motors. These directed movements were poleward (toward the nearest spindle pole, green arrowheads) and antipoleward (away from the nearest spindle pole, red arrowheads) on bipolar spindles (Fig. 2A-H). To more easily visualize individual or small clusters of Cut7-GFP, we used both photobleaching and photoactivation. Photobleaching near one spindle pole (marked by blue arrowhead near right SPB in fig. 2E) allowed additional observation of poleward and anti-poleward movement of Cut7-GFP near that SPB. Similarly, after activation of a subset of Cut7-PA-GFP (marked by blue arrowhead on left SPB in figure 2F), we were able to observe poleward and anti-poleward movement of Cut7-GFP.

**Fig. 2.**
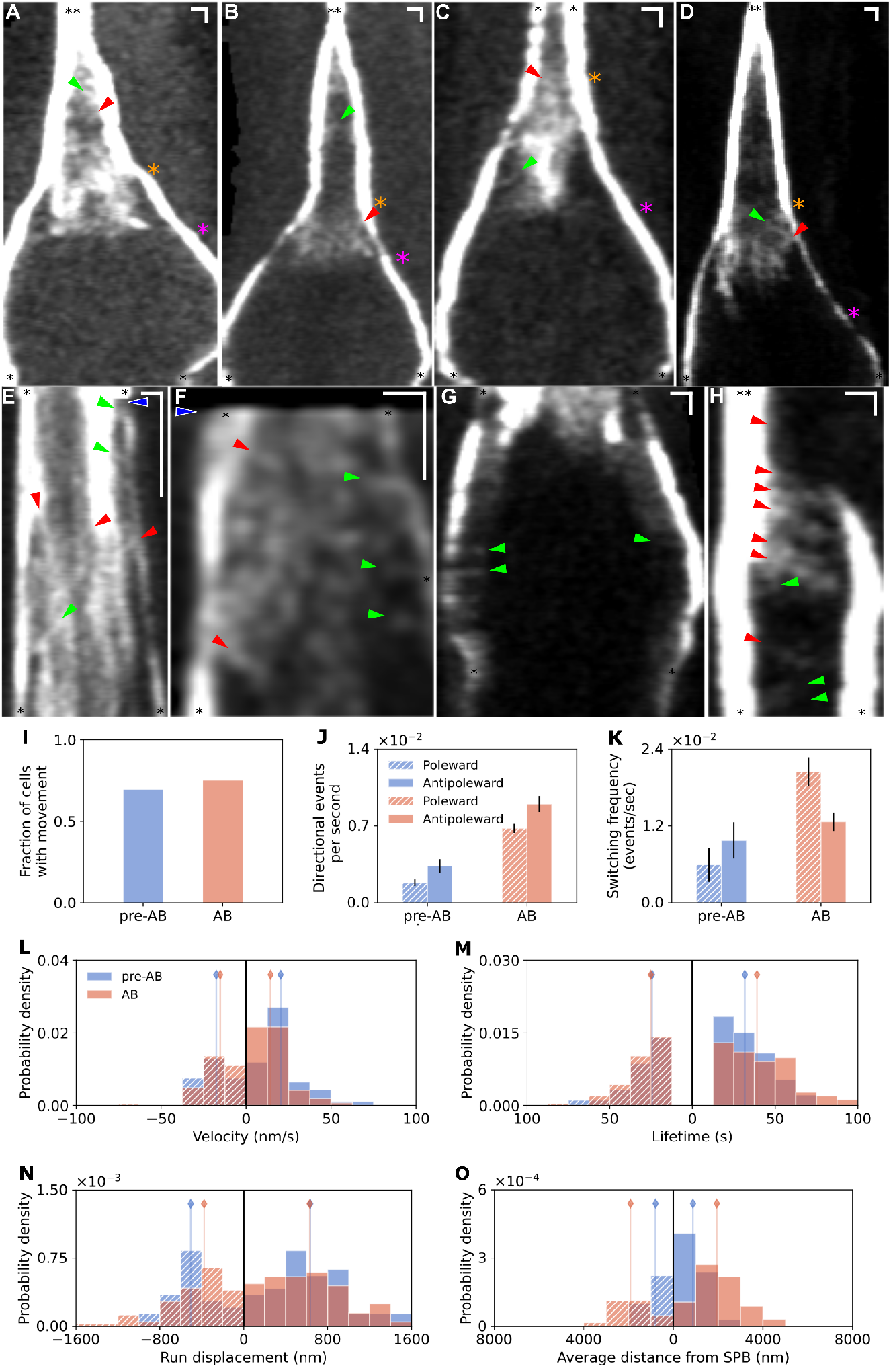
Localization and motility of Cut7p on bipolar spindles: changes during anaphase (A-D) Kymographs of Cut7-GFP movements on bipolar spindles from the onset of mitosis through the end of anaphase. Time reads from top to bottom. Orange arrowheads mark the transition from pre-anaphase B to anaphase B; magenta arrowheads mark the time when visible Cut7-GFP disappears from the interpolar spindle area yet remains at the spindle poles (E) A Fluorescence photobleaching experiment that reveals complex Cut7-GFP movements during anaphase B. The blue arrowhead marks the photobleaching event near the right spindle pole. Bright spots (clusters of Cut7-GFP) move more slowly than dim spots (likely small clusters or individual molecules). Near the photobleached pole, both poleward and anti-poleward movements are seen. (F) A Photoactivatable-GFP experiment showing complex Cut7-GFP movements during anaphase B. The blue arrowhead marks the time of photoactivation at the left SPB. Here, some fast moving clusters are visible. (G) A late anaphase B spindle showing pole-directed movements of Cut7-GFP that remains on the spindle. (H) Cut7-GFP localization during spindle pole separation at the onset of mitosis. Red arrowheads mark anti-poleward movements, green arrowheads mark movements toward the nearest SPB. Cut7-GFP moves from the left spindle pole and accumulates on the right spindle pole, though the final brightness of the right pole is great enough to suggest that Cut7-GFP may also move there from the nucleoplasm. SPBs: black stars; Vertical scale bars: 60 s; Horizontal scale bars: 1 *μ*m. (I – K) Comparison of Cut7-GFP motility before and after the onset of anaphase B. For quantification, see tables S1 and S2. (I) Fraction of cells in which visible movement occurred. (J) Frequency of visible directed movement. (K) Switching frequency out of each directed event. L-O) Quantification of cut7-GFP motility events. Values corresponding to minus-end directed movements are on the left side of the axis, plus-end directed movements on the right. Vertical bars represent median of the distribution. For quantification, see tables S1 and S2. (K) Velocity. (L) Lifetime. (M) Run displacement. (N) Average distance from the spindle pole including all points from each event.

We identified the onset of anaphase B as the time when the spindle elongation rate began to increase (orange stars, fig. 2A-D, Krüger et al. (2021)). Typically the intensity of Cut7-GFP spots near the spindle midzone was highest just after anaphase B onset. In early anaphase B, the kymographs suggested that Cut7p binding and/or movement increased, because a relatively bright, diffuse region of Cut7-GFP intensity appeared (fig. 2A-D, between orange and magenta asterisks). Subsequently, most Cut7-GFP intensity disappeared from the interpolar spindle (fig. 2A-D, magenta asterisks). After this time, any Cut7-GFP signal visible in the interpolar spindle typically moved toward the nearest spindle pole (Fig. 2G). The midzone of mid-to-late anaphase spindles contained little visible Cut7-GFP, consistent with previous findings that kinesin-6/Klp9 contributes more than Cut7p to anaphase B spindle elongation in fission yeast (Yukawa et al., 2018; 2019).

The *S. pombe* interpolar spindle in early mitosis includes MTs that extend from one pole all the way to the other (Ding et al., 1993;Ward et al., 2015). As a result, poleward and anti-poleward movement cannot be assigned to plus- or minus-end-directed movement. However, in late anaphase, MTs visible in spindle electron micrographs overlap only near the spindle midzone (Ding et al., 1993; Ward et al., 2015). Therefore, the poleward Cut7-GFP movement visible near the spindle poles in late anaphase (Fig. 2F-G) is likely minus-end-directed. This suggests that Cut7p can move toward the minus-ends of spindle MTs, consistent with its activity *in vitro* (Edamatsu, 2014; 2016; Britto et al., 2016).

We observed that Cut7-GFP was typically visible near one spindle pole earlier than the other. In *cut11-7* cells that have SPB insertion defects at restrictive temperature, we observed that even at permissive temperature, the asymmetry in early Cut7-GFP localization was exaggerated. Perhaps this is due to a delay in MT nucleation at the second SPB as observed previously for *cut11-6* (Zhang and Oliferenko, 2014). As a result, we observed bright Cut7-GFP signal near one spindle pole and little signal on the other, shortly after the poles separated (Fig. 2H). The kymograph shows that Cut7-GFP traveled primarily toward the dimmer spindle pole. In addition, these results suggest that Cut7-GFP moves in both directions between the spindle poles as the bipolar spindle assembles, and that Cut7-GFP signal can increase near a spindle pole due to movement from the interpolar spindle.

To further analyze Cut7p movement, we created custom Matlab software for kymograph analysis (Methods). We hand-traced visible Cut7-GFP tracks and then analyzed events (segments of tracks) that exhibited movement poleward or anti-poleward, or pauses (Fig. S2, S3). Additionally, we divided events into pre- and post-onset of anaphase B. For each movement event, we analyzed the velocity, lifetime, run displacement, and distance from the closest spindle pole. From the entire data set we identified the total number of cells that showed identifiable directed Cut7-GFP tracks, the frequency at which directional events occurred, and the state exit frequency (the switching frequency, defined as the rate at which a directional event ends by changing direction or pausing, Table S1, S2).

Consistent with visual inspection of the kymographs, Cut7p activity significantly increased after the onset of anaphase B (Fig. 2I-K). Before and after anaphase onset, the overall fraction of cells with bipolar spindles imaged that showed visible Cut7-GFP movement was similar, 69% pre-anaphase and 75% in anaphase (Fig. 2[I], Table S1). Despite this similarity, the number of directional events observed per minute was larger by a factor of 2-5 after anaphase onset (Fig. 2J). Both pre-anaphase B and in anaphase B, anti-poleward events occured slightly more frequently than poleward, consistent with the general tendency of Cut7-GFP to accumulate near spindle poles (Fig. 2J). Next, we measured the rate at which directed events ended, either due to pausing or changing direction (Fig. 2K). This rate increased slightly for anti-poleward events but by a factor of 3 for poleward events in anaphase B, meaning that direction switching and pausing occurred more frequently. Prior to anaphase B, poleward movement had a slightly lower state exit frequency, meaning that poleward movement persisted for longer than anti-poleward. During anaphase B, the state exit frequency of poleward movement was almost twice that of anti-poleward, meaning that anti-poleward movement lasted longer (was more persistent). Therefore, during early anaphase Cut7p movement events both occurred more frequently and switched direction or paused more frequently.

Analysis of individual directed movement events allowed us to determine motility parameters of poleward and anti-poleward events, both pre- and during anaphase B (Fig. 2[L-O], Table S2). Note that to facilitate comparison of poleward and anti-poleward movement, the left half of the graph/negative numbers show data for poleward movement, while the right half of the graph/positive numbers denote anti-poleward movement (always determined relative to the closest pole at the start of the event). The median velocity of Cut7-GFP movement was 15-20 nm/sec for both directions of movement and stages of mitosis (Fig. 2L). Therefore the change in Cut7p activity in anaphase does not appear to be driven by a change in motor velocity. The lifetime of directed events tended to be longer for anti-poleward movement (Fig. 2M). The run length was lower for poleward movement and decreased in anaphase B, while for anti-poleward events it was higher and stayed the same in anaphase B (Fig. 2N). By averaging all points from all tracks, we measured the average distance of visible tracks from the closest spindle pole and found that this increases by a factor of two in anaphase, reflecting a greater likelihood of Cut7 tracks to be visible farther from the spindle poles in anaphase B (Fig. 2O).

In summary, Cut7-GFP moved in both directions along bipolar spindles, and its direction-switching activity increased significantly in anaphase B. The speed of derected events was typically 15-20 nm/sec and lifetime 10-15 sec. Our observation of Cut7-GFP moving both poleward and antipoleward on bipolar spindles and switching direction was consistent with bidirectional motility on spindle MTs. However, interpolar spindle MTs are mixed in polarity, making it impossible to assign observed events as directed toward plus or minus ends of MTs in the diffraction-limited spindle. Motor movement near the poles in late anaphase B is likely plus-end-directed when anti-poleward and minus-end directed when poleward, but this is not definitive. To correlate Cut7p movement with MT polarity, we built on the observation that *S. pombe* monopolar spindles have minus-ends at the SPBs and plus-ends pointing outward. Therefore, we we sought to examine Cut7-GFP on monopolar mitotic spindles to determine whether the motor moves both toward plus and minus ends of microtubules.

### Cut7p moves bidirectionally on *cut11-7* monopolar spindles

Monopolar mitotic spindles can occur when a fission-yeast cell contains only one active SPB. These are observed reproducibly in fission yeast with temperature-sensitive mutations of *cut11* (West et al., 1998). In mitosis of these cells at restrictive temperature, a spindle MT bundle forms in which all the plus-ends are distal to the single pole (Fig. 3), allowing facile assignment of motor movement to polarity (Akera et al., 2015). We therefore tracked Cut7-GFP movement on this unipolar microtubule array to determine whether Cut7-GFP can move toward both the plus and minus ends of spindle MTs.

**Fig. 3.**
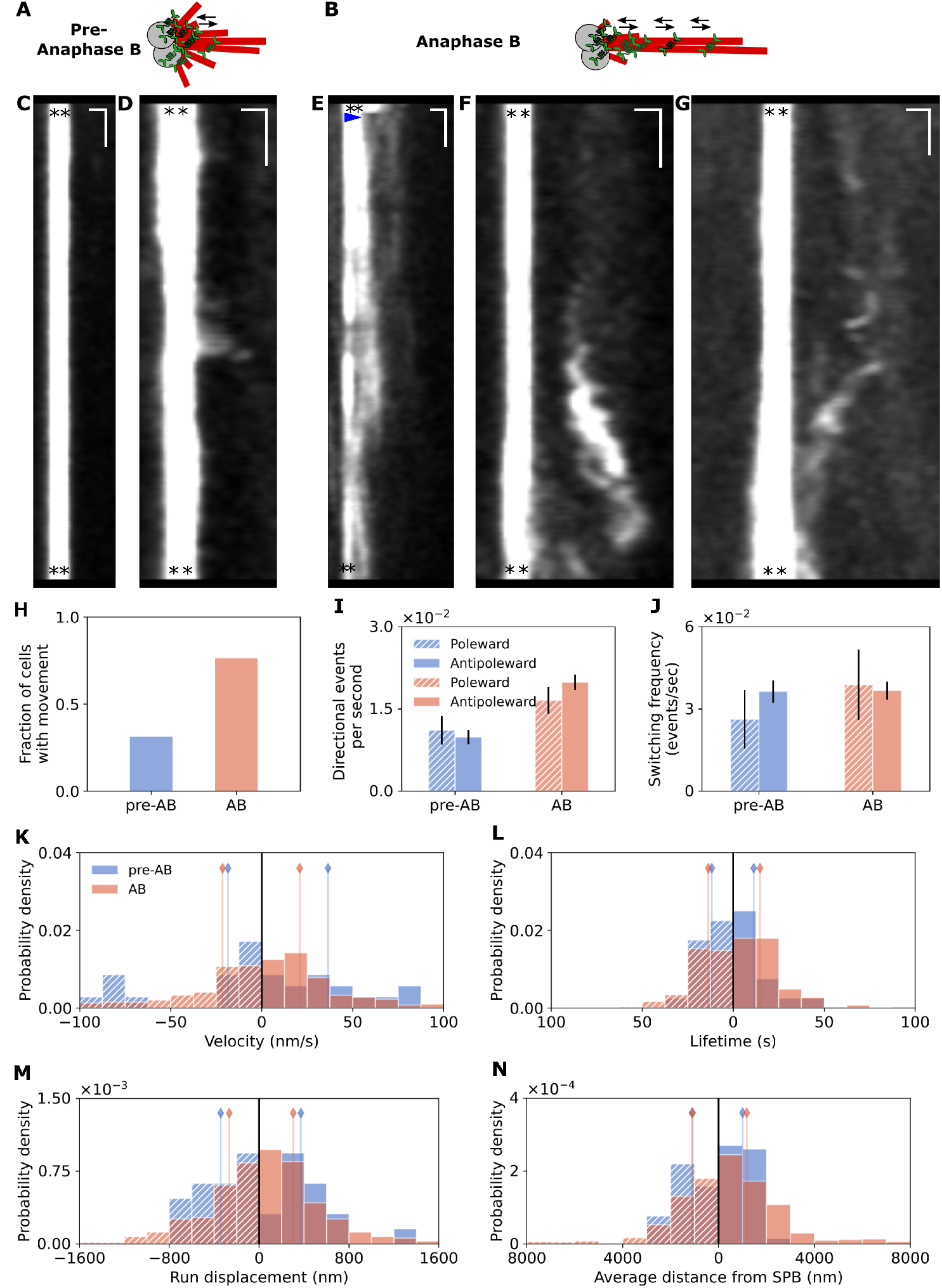
Localization and motility of Cut7p on monopolar spindles differs in short and long spindles (A-B) Cartoon of Cut7-GFP motor movements on monopolar spindles formed in cut11-7 cells held at a restrictive temperature for 2-4 hours before imaging. Spindles before and during anaphase B are compared. (C-G) Kymographs of Cut7-GFP motor movements on cut11-7 monopolar spindles. (C-D) movements pre-anaphase B. (E-G) movements during anaphase B. (E) is a photobleaching experiment; the blue arrowhead indicates the photobleaching event on the monopolar arm to the right of the spindle pole (asterisks). SPBs: black stars; Vertical scale bars: 30 s; Horizontal scale bars: 1 *μ*m. (I-O) Comparison of Cut7-GFP motility before and after the onset of anaphase B. For quantification, see tables S1 and S2. (I) Fraction of cells with any movements. (J) Frequency of visible directed movement. (K) Switching frequency out of each directed event. (L-O) Quantification of cut7-GFP motility events. Values corresponding to minus-end directed movements are on the left side of the axis, plus-end directed movements on the right. Vertical bars represent median of the distribution. For quantification, see tables S1 and S2. (L) Velocity. (M) Lifetime. (N) Run displacement. (O) Average distance from the SPB including all points from each event.

In *cut11-7* cells, monopolar spindles initially contained short MTs, similar to bipolar spindles in early mitosis (Fig. 3A). These short monopolar spindles contained relatively short, dynamic MTs. As time passes, some of the bundles elongated by almost a factor of two (Fig. 3B). Longer monopolar spindles contained one or more MT bundles that were more stable. The change from short to long monopolar spindles has been shown to occur at transition into an anaphase-B-like state, based on several cell-cycle markers (Krüger et al., 2021), despite the absence of chromosome segregation in these cells. Based on an average of 10 monopolar spindles that showed rapid elongation during our time of observation, we identified 2.4 *μ*m as the length at which this transition typically occurred, allowing us to define a boundary between spindles pre-anaphase and in anaphase B. This enabled us to compare the movement of Cut7-GFP on these two categories of monopolar spindle.

On all monopolar spindles Cut7-GFP localized primarily near the spindle pole, as observed previously (Akera et al., 2015). In this case, a kymograph of Cut7-GFP signal appears as a straight vertical bar (Fig. 3C). On the majority of pre-anaphase monopolar spindles, we observed no motion of Cut7-GFP. However in some cases these appeared to be short, dim events, likely corresponding to single motors or small clusters, characterized by movement a short distance from the SPB and then back (Fig. 3D). In longer anaphase monopolar spindles, brighter clusters were present either paused or moving along the spindle (Fig. 3E-G). In some cases we photobleached the signal near the SPB to allow visualization of smaller clusters (3E, blue arrow). On these longer monopolar spindles, we often observed Cut7-GFP along spindle MTs more distal to the SPBs (Fig. 3E-G). In some cases, Cut7-GFP moved in distinct tracks originating from the SPB (Fig. S4). For other events, the Cut7-GFP signal appeared brightly at one point along the spindle, without a clear track leading from the SPB (Fig. 3F-G). This may occur due to binding of motors directly from the nucleoplasm.

We quantified differences in Cut7-GFP movement on pre-anaphase and anaphase B monopolar spindles, and found that movement of Cut7-GFP away from the SPB occurred more frequently in anaphase B (76%) compared with pre-anaphase spindles (31%, fig. 3H, Table S1). Consistent with this, in anaphase B the frequency of observable directed events was a factor of 1.5-2 larger for both minus- and plus-end-directed movement (Fig. 3I). As a result, anaphase B spindles showed significantly more Cut7-GFP tracks than cells pre-anaphase B. The switching frequency out of poleward movement events appeared to increase upon entry to anaphase B (Fig. 3J), consistent with the increase in anti-poleward directed movement.

Quantitatively, directed events were similar in both directions during both pre-anaphase and anaphase. In addition, events were comparable to movement on bipolar spindles (Fig. 2). The median velocity increased slightly on monopolar spindles compared to bipolar (Fig. 3[K], Table S2), and directed events and run displacment were somewhat shorter as well sec (Fig. 3L,M). These changes were small but might reflect differences in Cut7p behavior during parallel versus antiparallel MT crosslinking, as has been shown for the *S. cerevisiae* kinesin-5/Cin8 (Gerson-Gurwitz et al., 2011). The average distance of Cut7-GFP from the SPB was ≈1 *μ*m (Fig. 3N). We found no significant correlation between Cut7-GFP spot intensity and particle velocity, lifetime, or run length. Together, these data provide evidence that Cut7p motility on the spindle can be toward either the plus or the minus end of MTs, and there is more motor activity in both directions as cells enter anaphase B. This adds additional evidence that this kinesin-5 moves toward minus-ends of spindle microtubules.

Cut7-GFP on monopolar spindles showed events moving in both directions, consistent with the idea that the motor moves bidirectionally. However, such bidirectional events could also occur if the motors can track dynamic MT ends. The median speed of directed Cut7-GFP tracks that we measured on bipolar and monopolar spindles was 15-20 nm s^−1^ (Fig. 2, 3, Table S2). This is slow compared to previous measurements of fission-yeast spindle MT dynamic instability, which found MT growth speed of 45-70 nm s^−1^ and MT shrinking speed of 60-110 nm s^−1^(Kalinina et al. (2012); Blackwell et al. (2017), Table S3), suggesting that the observed Cut7-GFP movement is unlikely to occur solely to microtubule plus-end tracking. However, it is possible that MTs with slower dynamics could be present in the spindle. Therefore, we perturbed both MT dynamics and Cut7p motility and examined the effects on Cut7-GFP motility on monopolar and bipolar spindles.

### Cut7p moves bidirectionally on *klp5*Δ monopolar spindles that contain more stable microtubules

To test whether the observed bidirectional Cut7p movement could be driven by spindle MT dynamics, we imaged Cut7-GFP on *cut11-7* monopolar spindles in cells containing deletions of kinesin-8/Klp5. Because Klp5p contributes to MT depolymerization, *klp5*Δ cells have microtubules and spindles that are 2-3 times longer and are more stable and bundled (West et al., 2001). In addition it has been shown that *klp5* deletion causes no change in the rates of microtubule growth and shrinkage; rather it decreases microtubule dynamicity, by reducing the frequency of both rescue and catastrophe (Unsworth et al., 2008). Consistent with previous work, we observed that monopolar spindles in *klp5*Δ *cut11-7* cells were significantly longer than those in *klp5*+ (Fig. 4A-G). As in *klp5*+ cells most Cut7-GFP on *klp5*Δ monopolar spindles remained near the SPB. If Cut7-GFP bidirectional movement occurs primarily due to MT dynamics, we would expect that Cut7p directional movement would occur less frequently and switch direction less frequently in *klp5*Δ cells.

**Fig. 4.**
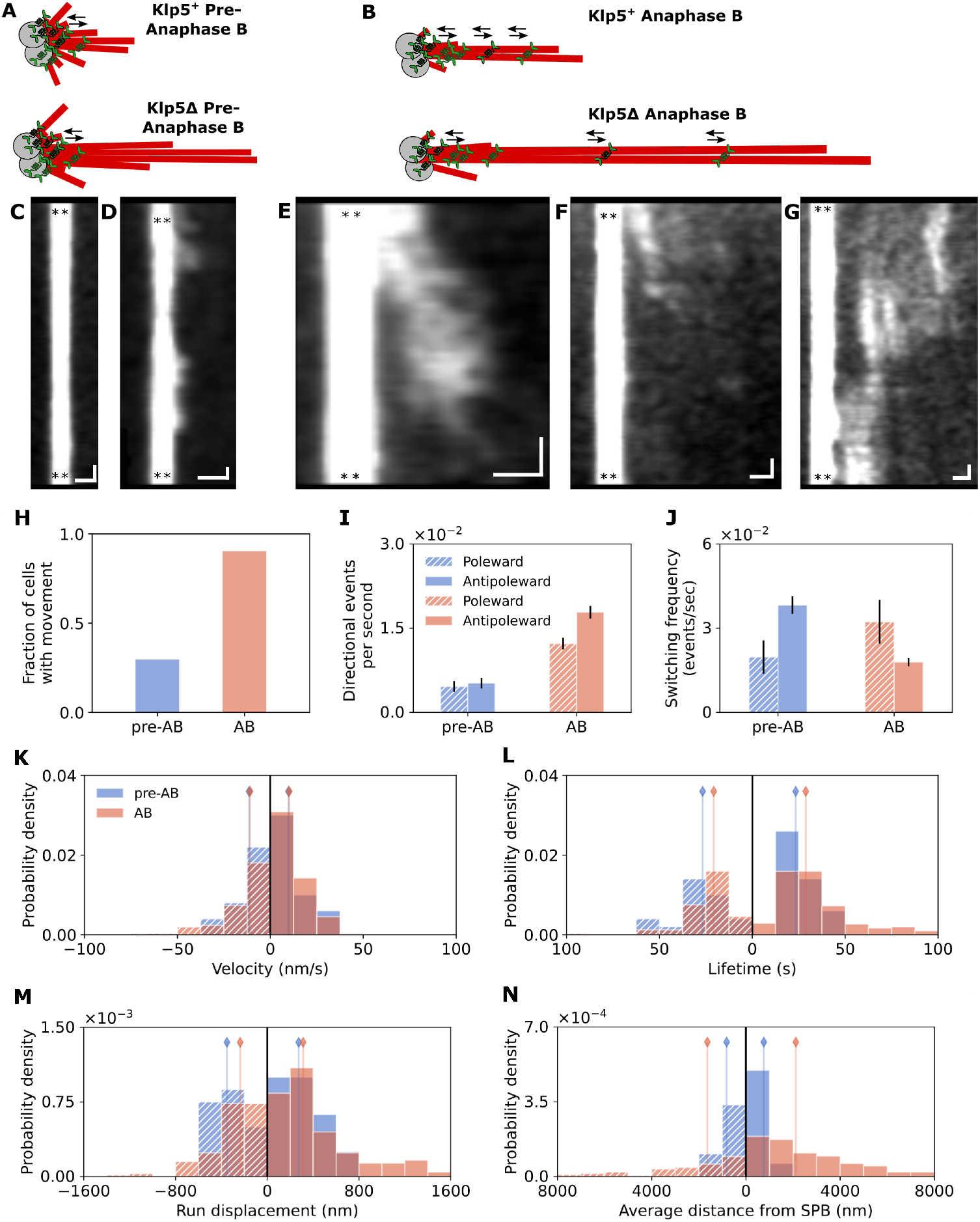
Longer monopolar spindles make the anaphase B increases in Cut7-GFP motility more evident. (A-B) Cartoon of Cut7-GFP motor movements on monopolar spindles in cut11-7 (upper) or in cut11-7, klp5Δ cells (lower), before and during anaphase B. (C-G) Kymographs of Cut7-GFP motor movements on klp5Δ, cut11-7 monopolar spindles. (C-D) show pre-anaphase B movements and (E-G) show anaphase B movements. SPBs: black stars; Vertical scale bars: 30 s; Horizontal scale bars: 1 *μ*m. (H-N) Comparison of Cut7-GFP motility on *klp5*Δ monopolar spindles before and after the onset of anaphase B. For quantification, see tables S1 and S2. (H) Fraction of cells with any movement. (I) Frequency of visible directed movement. (J) Switching frequency out of each directed event. (K-N) Quantification of cut7-GFP motility events. Values corresponding to minus-end directed movements are on the left side of the axis, plus-end directed movements on the right. Vertical bars represent median of the distribution. For quantification, see tables S1 and S2. (K) Velocity. (L) Lifetime. (M) Run displacement. (N) Average distance from the SPB including all points from each event.

We found that the fraction of pre-anaphase B cells that showed any spindle-associated Cut7-GFP motion was unaffected by the increased spindle length and was still only 30% (Fig. 4[H], Table S1) despite the longer length of spindle MTs on which events could take place. The fraction of cells with observable movement in *klp5*Δ cells increased during anaphase B to 90%. This fraction was higher in *klp5*Δ cells than in *klp5*+ cells (76%). This is the opposite of what would be predicted if MT dynamics drive the observed Cut7-GFP movements.

The number of observable directed Cut7-GFP events per minute was similar in the *klp5*+ and *klp5*Δ monopolar spindles (Fig. 4I), and the switching rates were comparable in the two strains (Fig. 4J, Table S1). This is consistent with the hypothesis that bidirectional motility arises from Cut7-GFP motility and the opposite of the prediction of the MT dynamics model. As in *klp5+* cells, motors in pre-anaphase B *klp5*Δ cells switched more frequently to minus-end directed movement, and switched more frequently to plus-end-directed movement in anaphase B (Fig. 4J). Overall, Cut7-GFP movement was relatively unaffected by the changing the length and dynamicity of monopolar spindle MTs (Fig. 4[K-N], Table S2). The median velocity of directed events decreased slightly compared to *klp5*+, but the velocity maintained a wide distribution (Fig. 4K). The lifetime increased slightly, again with a wide distribution (Fig. 4L). As a result the run length (Fig. 4M) was similar to measurements in *klp5*+ monopolar spindles. The average distance from the SPB increased in anaphase spindles, likely because the MTs were longer (Fig. 4N). Together, these data are consistent with the idea that that Cut7p moves bidirectionally on spindle MTs, rather than being moved solely by MT dynamics. To further test whether these movements are due to intrinsic Cut7p motility, we made several perturbations to Cut7p itself.

### Bidirectional motility is altered in Cut7 motor and tail mutants

To test whether perturbation of Cut7p can alter its movement on the spindle, we examined alterations to the motor that affect either the motor domain or the C-terminal tail. First, we asked whether Cut7p with no motor activity would change its localization on the spindle. If Cut7p movements are primarily driven by MT dynamics, we would expect to see a similar pattern of localization in motor-active Cut7p and motor-dead Cut7p. We therefore created a *cut7* mutant with a motor domain that could not bind ATP (*cut7-motor-dead*), constructed by changing three amino acids at residues 164-166. Previous work showed that this mutation blocks Cut7p movement but not microtubule binding (Akera et al., 2015). When *cut7-motor-dead* is the only allele of *cut7* in the cell, it is lethal. Therefore, we expressed it in a strain that also carries the temperature sensitive allele *cut7-446* (Hirano et al., 1986). Of the 25 cells with this genotype we imaged at permissive temperature (25 °C), 2 (8%) formed monopolar spindles and 23 (92%) formed bipolar spindles. Unlike Cut7-GFP (Fig. 1) and Cut7-446-GFP (Fig. 5A), Cut7-motor-dead-GFP was not enriched at spindle poles. Instead, it showed diffuse fluorescence along the entire length of the spindle (Fig. 5B). This is consistent with the idea that the typical spindle localization of Cut7-GFP depends on its minus-end-directed motility. In addition, this result suggests that Cut7-GFP accumulation at spindle poles does not occur solely due to microtubule dynamics.

**Fig. 5.**
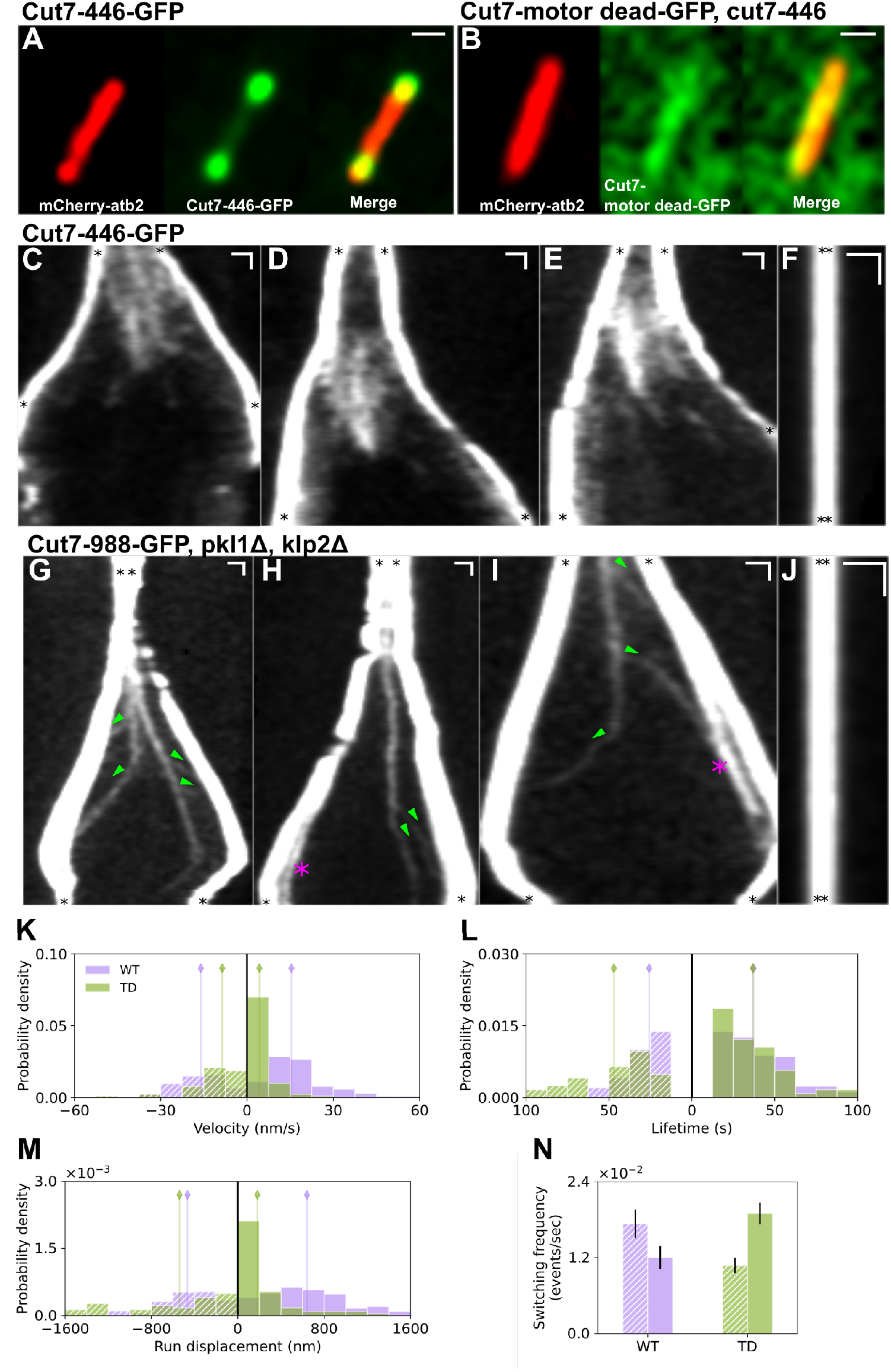
Mutations in Cut7 alter its motility on bipolar and monopolar spindles. (A-B) Two alleles of Cut7-GFP in cut7-446 cells at permissive temperature. (A) Dual color image of Cut7-446-GFP (green) on bipolar spindles, demonstrating sparse localization along the spindle MTs (red) but normal concentration at the poles (B) Analogous image of Cut7-motor dead-GFP in cut7-446 cells, showing some spindle binding but no concentration at the poles. (C-F) Kymographs of Cut7-446-GFP. Bipolar spindles (C-E) at permissive temperature show motility similar to cut7+ cells while monopolar spindles (F) at restrictive temperature show severely reduced plus end directed motility compared to Cut7+-GFP. (G-M) Cut7-tailΔ-GFP kymographs and data. Removal of the cut7 tail domain in the presence of pkl1Δ and klpΔ leads to reduced motility in bipolar spindles (G-I) and reduced plus end directed motility in monopolar spindles (J). SPBs: black stars; Vertical scale bars: 60 s; Horizontal scale bars 1 *μ*m. (K-N) Comparison of Cut7-GFP and Cut7-988-GFP motility Values corresponding to poleward directed movements are on the left side of the axis, anti-poleward directed movements on the right. Vertical bars represent median of the distribution. For quantification, see table S2. (K) Velocity. (L) Lifetime. (M) Run displacement (N) Switching frequency out of each directed event.

Next, we sought to determine whether the *cut7-446* temperature-sensitive mutation affects Cut7p motility. This is a single point mutation, I954T, near to the start of the tail region (Yukawa et al., 2018). As mentioned above, at permissive temperature these cells formed bipolar spindles with Cut7-446-GFP localization similar to Cut7-GFP (Fig. 5A). Further, kymographs of Cut7-446-GFP appeared similar to those of Cut7-GFP throughout mitosis (Fig. 5C-E). We then shifted *cut7-446* cells to restrictive temperature (37 °C), which leads to temperature-sensitive lethality by blocking normal SPB separation (Hagan and Yanagida, 1992). In 98% of the monopolar spindles that formed (N=104), Cut7-446-GFP was visible only near the SPBs, with no directed movement events. This suggests that at restrictive temperature the motor either cannot move or only retains minus-end-directed motion; we observed almost no plus-end-directed motion (Fig. 5F), in contrast to *cut11-7* monopolar spindles in which Cut7-GFP showed plus end-directed movements in 36% of spindles (Fig. 3H). This observation is consistent with Cut7p immunofluorescent detection in *cut7-446* cells (Hagan and Yanagida, 1992). This suggests that movement of Cut7-GFP is due to its intrinsic motility, and not primarily by tracking plus-ends of dynamic MTs. In addition, this single amino acid change appeared to disrupt plus-end movement at restrictive temperature.

To further perturb Cut7, we truncated the C-terminal tail. The C-terminal tail contains the conserved BimC box containing a Cdk phosphorylation site (Drummond and Hagan, 1998), and point mutations in this region lead to alleles that are temperature sensitive for growth (Rodriguez et al., 2008; Akera et al., 2015). Further, tail truncation in other kinesin-5s can impair MT crosslinking and sliding (Hildebrandt et al., 2006; Bodrug et al., 2020). Therefore we asked whether truncation of the tail at the end of the predicted coiled-coil domain (Fig. S5) affects Cut7p motility. The truncation we constructed removed 96 amino acids from the tail beyond amino acid 988. This mutation appeared to be lethal, as we were unable to produce transformants with *cut7-988* as the sole *cut7* allele in the cell. This is consistent with observations in budding yeast that deletion of the kinesin-5/Cin8 tail is lethal in the absence of kinesin-5/Kip1 (Hildebrandt et al., 2006). To allow observation of motility of Cut7-988-GFP, we examined its phenotype in cells with deletions of *pkl1* and *klp2*, genes that encode two minus end-directed kinesin-14s. Because Pkl1p and Klp2p are antagonistic to Cut7, deleting them allows study of lethal alleles of *cut7* (Pidoux et al., 1996; Troxell et al., 2001; Rincon et al., 2017; Yukawa et al., 2018; Lamson et al., 2020;Yukawa et al., 2020). This combination of mutations produced viable cells.

On bipolar spindles, Cut7-988-GFP localized predominantly to the spindle poles, similarly to Cut7-GFP (Fig. 5G-I). This suggests that the motor has sufficient activity to move to or bind to microtubule minus ends, in contrast to Cut7-motor-dead-GFP (Fig. 5B). However, Cut7-988-GFP appeared present at lower levels on the interpolar spindle than Cut7-GFP in the same strain background (Fig. 1E-F compared to Fig. 5G-I). This suggests that the Cut7 tail may play a role in either plus-end-directed movement or in stabilizing microtubule binding once the motor has reached the midzone. The Cut7-988-GFP that was observed near the midzone in early mitosis lay in faint clusters with reduced movement (Fig. 5G-I). In late anaphase B, however, similar clusters showed processive movement toward spindle poles (Fig. 5G-I, green arrowheads). In several cases we observed fluorescent trajectories in the kymographs splitting in two. In some cases, the sum of the average brightness of two individual tracks equaled the brightness of the initial track (Fig. 5G,H), suggesting that the initial track may have been two motor tetramers that then separated into single-motor events. The poleward movement typically continued until the signal reached the spindle pole. On monopolar spindles, we observed no plus-end-directed movement (Fig. 5J), consistent with the lethal phenotype of this deletion allele when kinesin-14s are present. Together, these data suggest that Cut7-988-GFP has lost most plus end-directed motility but retains minus end-directed motion.

To test whether the minus-end-directed movement we observed in late anaphase was due to Cut7-988-GFP motor activity or to immobile Cut7p on depolymerizing microtubules in the final stages of anaphase, we monitored the time at which the mCherry-tubulin signal in the spindle decreased and the time when Cut7-988-GFP moved poleward. The Cut7p poleward movement typically preceded the microtubule signal decrease (Fig. S6). Thus, late anaphase Cut7-988-GFP movement is likely minus-end-directed motor activity. At the same stage of anaphase, we also observed bright, fluorescent, relatively immobile clusters of Cut7-988-GFP near the spindle poles (Fig. 5H,I, magenta asterisks).

We quantified Cut7-988-GFP movement and compared it to Cut7-GFP movement on bipolar spindles. Because *cut7-988-GFP* cells appeared to show defects in spindle elongation, we could not easily identify the onset of anaphase B from spindle length changes. Therefore, we examined all directed movement events, independent of stage of mitosis, for both Cut7-GFP and Cut7-988-GFP (Fig. 5[K-N], S6, Table S1, S2). The speed of Cut7-988-GFP movement was 2-3 times slower than Cut7-GFP, depending on the direction of movement (Fig. 5K). This is in contrast to *Drosophila melanogaster* kinesin-5/KLP61F, which showed faster motility *in vitro* when the tail was truncated (Bodrug et al., 2020). The lifetime of poleward movement events was nearly two times longer for Cut7-988-GFP than for Cut7-GFP, while the lifetime of anti-poleward movement was comparable (Fig. 5L). As a result, the run displacement of anti-poleward movement was ~ 3 times smaller for Cut7-988-GFP than for Cut7-GFP (Fig. 5M). Consistent with this, the switching rate out of anti-poleward movement was higher than the exit rate from poleward movement for Cut7-988-GFP; this is the reverse of the trend for Cut7-GFP (Fig. 5N). This is again consistent with the hypothesis that Cut7p bidirectional movement is a result of motor activity, and while MT dynamics may affect its motility they are not the primary driver. Further, our results show that the C-terminal tail beyond amino acid 988 is a determinant of Cut7 motor activity, with a particular contribution to plus-end-directed movement.

### Human Eg5 that replaces Cut7p shows altered localization and motility on fission-yeast spindles

While several mutations to *cut7* appeared to either abolish its movement or favor MT minus-end-directed motility (Fig. 5), we did not identify *cut7* mutants with a greater propensity for plus-end-directed movement. An alternative to study plus-end-directed movement is the human kinesin-5/Eg5, a plus-end-directed motor that has recently been shown to complement *cut7* as the sole kinesin-5 in *S. pombe* (Hwang et al., 2022). Consistent with prior work, we found that cells containing Cut7-GFP or Eg5-GFP expressed at low levels in the *cut7* deletion background were viable (Fig. 6, S7). While Cut7-GFP localized brightly to the spindle poles and more dimly along spindle MTs (Fig. 6A), Eg5-GFP was visible along the spindle with little to no enhancement at the poles (Fig. 6B). This is consistent with our other results suggesting that the bidirectional motility of Cut7 contributes to its pole localization: plus-end-directed Eg5 would therefore not be predicted to accumulate at the spindle poles.

**Fig. 6.**
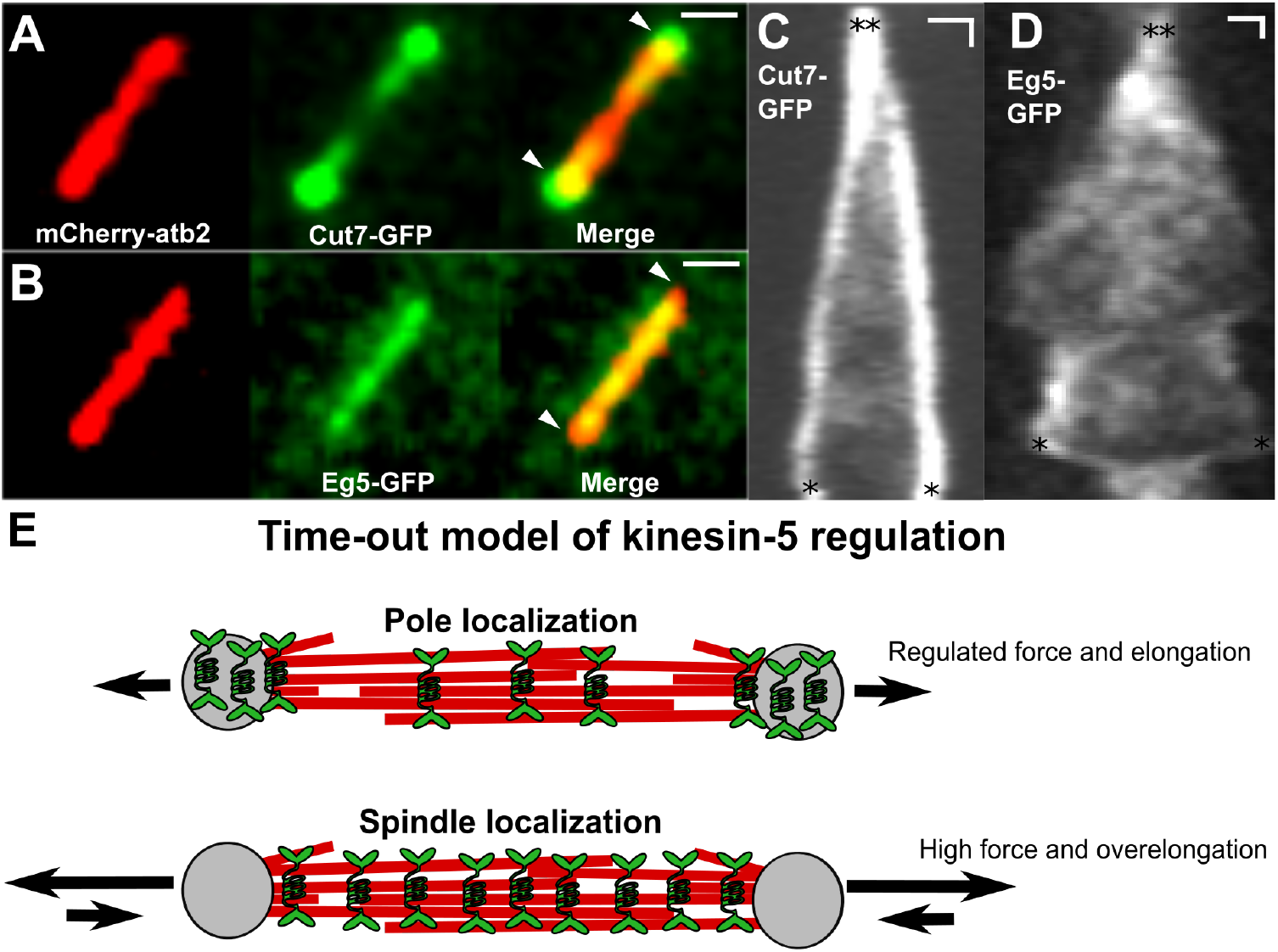
Eg5-GFP shows altered localization on bipolar spindles. (A-B) Dual color images of bipolar spindles. (A) Cut7-GFP (green) showing typical enrichment at the SPBs (arrowheads) and binding along the MTs (red). (B) SPB enrichment in Eg5-GFP (green) is no longer present (arrowheads) and Eg5-GFP binds uniformly along the spindle. (C-D) Kymographs of bipolar spindles showing early movements after the onset of mitosis. (C) Cut7-GFP shows consistent SPB enrichment and movements along the interpolar spindle. (D) Eg5-GFP shows reduced SPB enrichment and more uniform binding along the spindle. Spindle shortening events are also observed. SPBs: black stars; Vertical scale bars: 60 s; Horizontal scale bars: 1 *μ*m. (E) Cartoon model demonstrating the hypothesis that motor localization on the spindle affects separation force and spindle elongation.

To further examine the dynamics of motor localization over time, we compared kymographs of Cut7-GFP and Eg5-GFP on bipolar spindles. The pole localization is noticeable for Cut7-GFP (Fig. 6C), while Eg5-GFP is visible at the spindle poles only rarely and transiently (Fig. 6D). We also observed that in Eg5-GFP cells, initial spindle pole separation often occurred more quickly and was followed by spindle shortening events in about 20% of cells (Fig. 6D). These results suggest that when a purely plus-end-directed kinesin-5 motor is present on the fission yeast spindle, it may be over-active and drive premature spindle elongation followed by shortening.

## DISCUSSION

### Cut7-GFP moves bidirectionally on spindle microtubules with regulated motility

Our results show that Cut7p can move in cells toward both plus- and minus-ends of mitotic microtubules, similar to its bidirectional motility *in vitro* (Edamatsu, 2014; 2016; Britto et al., 2016). After verifying that Cut7-GFP localized on the spindle, as observed in previous work (Fig. 1, Hagan and Yanagida (1992); Drummond and Hagan (1998); Yukawa et al. (2015)), we examined the directional movement of Cut7-GFP on both bipolar and monopolar spindles. Cut7-GFP fluorescent signal moved in both directions along bipolar spindles (Fig. 2), which is consistent with bidirectional motility but does not definitively demonstrate it because the spindle contains extensive overlapping antiparallel microtubules. The bidirectional movement also occurred on monopolar spindles that contain microtubules of only one orientation (Fig. 3, 4). We note that Cut7-GFP movement on bipolar and monopolar spindles was similar, and occurred during comparable time in mitosis in all cells. The similar motility observed therefore suggests that motor direction is defined by features of the motors themselves, rather than being strongly influenced by microtubule polarity and parallel versus antiparallel crosslinking. Consistent with this view, we observed that individual fluorescent spots can change their direction of motion [Fig. 3D-G].

The speed of directed events that we measured was ~ 15-30 nm s^−1^ for both directions of movement (Fig. 2, 3, Table S2). In previous experiments with purified Cut7, surface-bound motors drove microtubule motion in gliding assays at similar speeds, ~ 10-30 nm s^−1^ (Edamatsu, 2014; Britto et al., 2016), although the speed was higher (~ 200 nm pers) for minus-end-directed movement at low concentration (Britto et al., 2016). Motion of Cut7 on single microtubules occurred at 150-200 nm s^−1^(Edamatsu, 2016). Thus our velocity measurements are consistent with the slower velocities measured *in vitro* and the speed of KLP61F *in vitro* (Bodrug et al., 2020).

Our findings suggest that while Cut7p movement has a stochastic component, it is regulated and changes as mitosis progresses. In particular, we found that the activity of Cut7-GFP increased at anaphase B onset on both bipolar and monopolar spindles. We observed a larger percentage of cells that showed Cut7-GFP movement, a greater frequency of directed events, and an increased rate of switching in anaphase B (Fig. 2, 3, 4, table S1).

These observations of increased Cut7-GFP activity and fluorescence signal in the spindle midzone is consistent with its contribution to anaphase spindle elongation (Yukawa et al. (2019), Fig. 2). The fact that most Cut7-GFP signal disappears from the spindle midzone not long after anaphase B starts is, however, consistent with previous work showing that kinesin-6/Klp9 is the primary motor driving spindle elongation in *S. pombe* (Fu et al., 2009). By late anaphase, the Cut7-GFP still visible on the spindle appears to be primarily minus end-directed, and thus likely does not contribute significantly to spindle elongation (Fig. 2, 5). Therefore, it is likely that any contribution of Cut7p to spindle elongation by antiparallel microtubule sliding likely occurs early in anaphase.

### Motor and tail domains both contribute to Cut7 bidirectional motility

We find that both the motor and C-terminal tail domains of Cut7 contribute to its bidirectional movement and localization. The mutations we examined appeared to limit the motility of Cut7p toward MT plus ends, which suggests that the ability of Cut7p to move toward MT plus ends is more fragile that its minus-end-directed motion. Both a point mutation in the tail region (*cut7-446*) and a tail deletion (*cut7-988*) favor the motor’s pole localization and reduce its plus-end-directed motility (Fig. 5). The results also point to a role of the C-terminal tail domain as a site for motor regulation. Some properties or modification of the tail may be important for inducing plus end-directed motion in Cut7p.

While wildtype *S. cerevisiae* Cin8 localizes to the minus ends at the SPB of a pre-assembled (monopolar) spindle, a mutant lacking a C-terminal tail (but still containing a nuclear localization signal) localizes to the plus ends of the monopolar spindle (Shapira et al.,2017). On a bipolar spindle, this mutant localizes similarly to a wildtype Cin8, at the spindle poles and the interpolar spindle (Düselder et al., 2015). Therefore, the tail truncation of Cin8 appears to favor its plus-end-directed motility. In contrast, here we show that the Cut7-988-GFP tail deletion mutant compared to full-length shows an increased bias toward the minus ends of both bipolar and monopolar spindles.

Truncation of the *Drosophila melanogaster* kinesin-5/KLP61F tail causes it to move more quickly and cluster less *in vitro*, possibly because tail-motor interactions alter the motor conformation to slow ATP hydrolysis (Bodrug et al., 2020). In contrast, we find in cells that the tail-truncated Cut7-988-GFP moves more slowly; however, we also see decreased clustering. Despite these differences, both motors appear to be impaired in sliding MTs. For KLP61F decreased MT sliding force generation was directly measured *in vitro* (Bodrug et al., 2020), while our results indirectly suggest that Cut7-988p has defects in force generation because it is lethal unless the opposing kinesin-14s are also deleted.

The bidirectional movement of Cut7p that we observed could in principle be driven by tip-associated Cut7p on dynamic spindle MTs. To test this idea, we examined how Cut7-GFP localization and motility changed in cells when MT dynamics or Cut7p motility were perturbed. Additionally, we compared localization of Cut7p to human Eg5. Deletion of kinesin-8/Klp5 induces elongation of cellular microtubules and decreases their dynamicity (Unsworth et al., 2008). We found that Cut7-GFP motility did not significantly change on monopolar spindles in *klp5*Δ cells (Fig. 4), supporting that idea that microtubule dynamics are not the primary driver of the observed Cut7-GFP movement.

In contrast, perturbations to Cut7p itself did lead to changes in its localization and motility (Fig. 5). Our observation that Cut7p accumulates poorly at SPBs when its ATPase activity is eliminated (Fig. 5B) provides evidence that Cut7p spindle pole accumulation is at least in part motor-driven. Consistent with this idea, plus-end-directed human Eg5 can replace Cut7p in *S. pombe* (Hwang et al. (2022), Fig. S7), but does not strongly localize to spindle poles (Fig. 6B). We conclude that Cut7p motor activity contributes significantly to the directed movement we have described.

### Function of fission-yeast versus budding-yeast kinesin-5s

The budding yeast *S. cerevisiae* contains two kinesin-5s, Kip1 and Cin8. Of these, Cin8 appears to be more similar to Cut7 (51.2% BLAST sequence identity versus 35.4% sequence identity with Kip1). Both budding yeast motors are strongly localized to the spindle poles and more weakly at or near the spindle midzone, but neither of them accumulates there until some time into anaphase (Gerson-Gurwitz et al., 2011; Fridman et al., 2013). This pattern of localization is somewhat similar to that of of Cut7p in *S. pombe*. However, Cut7p does not concentrate at the midzone in late anaphase, as Cin8p does, probably reflecting the fact that kinesin-6/Klp9 is important for late anaphase B spindle elongation in fission yeast, while the kinesin-5s play this role in budding yeast. Both Cin8 and Cut7p have been shown to be bidirectional *in vitro* (Roostalu et al., 2011; Gerson-Gurwitz et al., 2011; Edamatsu, 2014; 2016; Britto et al., 2016), and Cin8 also moves bidirectionality on spindle MTs *in vivo* (Gerson-Gurwitz et al., 2011; Fridman et al., 2013). Our work thus confirms that Cut7p moves bidirectionally on the spindle *in vivo* as well. Both motors are important for separation of SPBs to establish a bipolar spindle (Hoyt et al., 1992; Roof et al., 1992; Hagan and Yanagida, 1990; 1992), suggesting that both exert plus-end-directed forces early in mitosis. In contrast to Cut7p, Cin8 appears to persist in this role throughout mitosis, while Cut7p passes that role on to a different motor.

In *S. cerevisiae*, distinct, processive Cin8 tracks appear common (Gerson-Gurwitz et al., 2011). In *S. pombe*, clear tracks can be identified but appear less common (Fig. 2). Instead, a region of dim Cut7-GFP fluorescence along the spindle, perhaps composed of many moving motors, is more commonly observed. We observed distinct tracks occasionally in cells with Cut7-GFP and frequently in cells with the tail truncated Cut7-988-GFP (Fig. 5). Therefore, the region of dim Cut7-GFP fluorescence on bipolar spindles likely reflects complex movement of many motors/clusters.

The minus-end-directed motility of Cin8 depends on phosphorylation in the motor domain on Cdk1 sites not conserved in *cut7* (Shapira and Gheber, 2016). By contrast, we find that the Cut7p perturbations we examined decreased or abolished plus-end-directed movement. Our results therefore suggest that Cut7p is more intrinsically minus-end-directed and acquires plus-end directionality by some mechanism that has not yet been clearly identified. This may be due to motor crowding (Britto et al., 2016), phosphorylation analogous to the regulation of Cin8, or some other mechanism.

### Function of Cut7p bidirectional motility and pole localization

It is well established that kinesin-5s are able to crosslink and slide antiparallel MTs to separate spindle poles, a function which depends on plus-end-directed motility of kinesin-5 tetramers. Our work further supports the importance of kinesin-5 plus-end-directed motility, because *cut7* mutations that eliminate motility or reduce plus-end-directed events either abolish or impair spindle-pole separation (Fig. 5). Given the importance of plus-end-directed motility, the importance of bidirectional motility/trafficking and pole localization has remained unclear.

In previous work, we used a computational model of *S. pombe* spindle assembly to propose that localization of kinesin-5/Cut7 near the spindle poles may be required for proper spindle-pole separation (Blackwell et al., 2017; Edelmaier et al., 2020). This was based on model simulations in which a processive, purely plus-end-directed kinesin-5 localized far from the SPBs in monopolar spindles and failed to separate spindle poles. Experimental work on budding-yeast Cin8 reached a similar conclusion (Shapira et al., 2017). However, the experimental results we present here appear to rule out that hypothesis. Importantly, human Eg5 can replace *cut7* and assemble a bipolar spindle (Hwang et al., 2022) despite being a plus-end-directed kinesin-5 and not significantly localizing to *S. pombe* spindle poles (Fig. 6). Therefore, minus-end-directed motility is not required for bipolar spindle assembly in fission yeast. How to reconcile this with the computational modeling results is an interesting question for future work. We speculate that perhaps Eg5 has higher turnover that allows it to unbind and avoid becoming localized primarily at the plus-ends of spindle microtubules in early mitosis.

We observed that replacement of Cut7p with low-level expression of Eg5 in *S. pombe* can lead to faster initial separation of spindle poles, a longer spindle, and transient spindle shortening (Fig. 6D). Additionally, recent work found that high-level expression of either human Eg5 or Cut7p in *S. pombe* led to over-elongated spindles and was lethal (Hwang et al., 2022). Together, these results suggest that a highly expressed or purely plus-end-directed kinesin-5 that localizes along spindle MTs may produce excessive sliding force that over-elongates the spindle. Therefore, we propose a new hypothesis that spindle-pole localization of kinesin-5 may sequester the motor in a “time-out” at a location where its sliding activity is low (Fig. 6E). In this view, the spindle-pole localization may decrease motor activity. This model will be of interest to test in future work.

## MATERIALS AND METHODS

### Strain construction by crossing

Cells were cultured using standard techniques (Moreno et al., 1991). Existing strains (table S4) were crossed and the desired phenotype isolated using random spore analysis. The *cut11-7* and *cut7-446* mutations were identified by replica plating colonies from the spore analysis onto YE5S plus phloxin B agar plates. After 2-3 d at 37°C, dark-pink colonies were identified as positive for the mutations. All other genes were identified using EMM plates without the relevant supplement for auxotrophic mutants or YE5S plates with the relevant antibiotic. The fluorescent label for microtubules was obtained by expressing an mCherry-α-tubulin-chimera at a low level (~10 percent wild type α-tubulin), as used previously (Yamagishi et al., 2012; Gergely et al., 2016; Blackwell et al., 2017). This low-level labeling reduces possible tag-related perturbations to microtubule dynamics. Cells with kinesin-5 mutation or deletion are susceptible to chromosome missegregation and other mitotic abnormalities. To avoid problems resulting from these abnormalities, cells for each experiment were stored at 4°C and reisolated from frozen stocks every 2-4 weeks.

### Plasmid and strain construction by molecular biology

Oligonucleotide primers were purchased from Integrated DNA Technologies (Coralville, IA). Restriction enzymes and Phusion HF DNA polymerase were purchased from New England Biolabs (Ipswich, MA). DNA was prepared using Qiaprep Spin Miniprep Kit and polymerase chain reaction (PCR) products were purified using Qiaquick PCR Purification Kit, both from Qiagen (Germantown, MD). *S. pombe* genomic DNA was prepared using YeaStar Genomic DNA Kit from Zymo Research (Irvine, CA). DNA sequencing was performed by Quintarabio (Hayward, CA). DNA concentration was determined using a Thermo Scientific Nanodrop 2000. Strains were verified by PCR and sequence analysis of genomic DNA. At least two different transformant strains were analyzed in experiments.

### Strain construction for Cut7-photoactivatable(PA)-GFP

PA-sfGFP (Addgene 54579) was amplified by PCR using oligonucleotide primers Pac1-paGFP-F (GGGTTAATTAACGTGAG-CAAGGGCGAGGAG) and Asc1-paGFP-R (AGTGGCGCGCCCTACTTGTACAGCTCGTCCATGCC) and the product was cloned into pFA6a-GFP(S65T)-kanmx6 (Addgene 39292) digested with Pac1/Asc1. The resulting plasmid was amplified by PCR using oligonucleotide primers Cut7-Cterm-pFA6a-GFP/paGFP-F (AATTCATCAAGAACTAGTCTTTTGCGGAGTAGCAGAAGT-GCCTATTCCAAAATGAAACGACGGATCCCCGGGTTAATTAA) and Cut7-Cterm-pFA6a-GFP/paGFP-R (GATGTAATACATTTC-TATTGTATTTTCGTCCATTAAGTATAAATTAATAAAATCGTGTCAGAATTCGAGCTCGTTTAAAC) and the product was used to transform *S. pombe* using a lithium acetate method (Okazaki et al., 1990).

### Strain construction for Cut7-446-GFP and Cut7-988-GFP

For *cut7-446-GFP*, strain MB921 was transformed with PCR product obtained using plasmid pFA6a-GFP(S65T)-kanmx6 (Addgene 39292) and oligonucleotide primers Cut7-Cterm-pFA6a-GFP/paGFP-F and Cut7-Cterm-pFA6a-GFP/paGFP-R. For *Cut7-988-GFP*, strain MB1147 was transformed with PCR product obtained using plasmid pFA6a-GFP(S65T)-kanmx6 (Addgene 39292) and oligonucleotide primers Cut7-988-Cterm-pFA6a-GFP-F (TTAAAGGGAACGACATCACTTGCTAATCATACTAATGAATTACTTGGTTTAG-GAGATGAACGGATCCCCGGGTTAATTAA) and Cut7-Cterm-pFA6a-GFP/paGFP-R. The choice to truncate at amino acid 988 was based on coiled-coil prediction (Lupas et al. (1991), Fig. S5).

### Strains expressed from heterologous promoter

The genes for *cut7*, *cut7-motor-dead*, and Eg5 were cloned using PCR into plasmid 462 (Addgene 89065) containing a Z3EV promoter (Ohira et al., 2017) modified to contain Asc1/Srf1 sites just upstream of the GFP gene. The motor dead mutation (G164A-K165A-T166A) was cloned by PCR from strain PB951 (MB947, Akera et al. (2015)) into the 462-cut7 plasmid. These plasmids were cut at Aatll and integrated at the *his7* gene in Strain FY31411 (yFS949, MB1138). The strain was crossed to produce strain MB1164. Estradiol was not used in our experiments; rather *cut7* or Eg5 were expressed at a constitutive lower level.

### Live-cell imaging

All microscopy images and related datasets were obtained from confocal microscopy using live-cell preparation. Cells were grown at 25°C on YE5S plates for 48 hours and restreaked every 12 hours. A small volume of exponentially growing cells was removed from the petri dish and placed in 10 μL of EMM. EMM was filtered with a 0.2 *μ*M cellulose acetate filter to reduce background fluorescence. EMM was placed onto 22 × 60-mm coverslips coated with 8 *μ*L of lectins from *Bandeiraea simplicifolia* (Sigma-Aldrich). These coverslips were pre-equilibrated to the appropriate temperature of either 25 or 37°C. Bipolar spindles were imaged at 25°C and monopolar spindles were imaged at 37°C. To obtain sufficient monopolar spindles, *cut11-7* cells were placed at 37°C for 2-4 hours and then imaged at this restrictive temperature. Cells were transferred from a 37°C incubator to the pre-warmed microscope in less than 30 seconds to prevent monopolar spindles cooling down and becoming bipolar. Temperature was maintained with ±0.1°C precision using a CherryTemp temperature controller (Cherry Biotech, Rennes, France). Spinning-disk confocal microscopy was performed on a Nikon Eclipse Ti microscope described previously (Gergely et al., 2016; Blackwell et al., 2017; Edelmaier et al., 2020) . Time-lapse image stacks were obtained using the EM gain laser settings on the Nikon illumination system and number of Z-planes described previously (Edelmaier et al., 2020).

### PA-GFP and fluorescence photobleaching experiments

PA-GFP experiments used a 405 nm, 50 mW coherent Obis laser (Santa Clara, CA). Before photoactivation, 4 image stacks with illumination from the 561 nm and 488 nm lasers were obtained to record the location of the microtubules and confirm that no GFP signal existed prior to activation. Photoactivation was then performed on either a monopolar microtubule bundle or near the spindle pole with the laser at 10% power and an exposure time between 15-1000 ms. After photoactivation, time-lapse image stacks were obtained continuously using the 488 nm illumination laser. For fluorescence photobleaching experiments, the same 405 nm laser was used. Before photobleaching, 4 image stacks with illumination from the 488 nm laser were obtained to record the location of the spindle poles and Cut7-GFP. Photobleaching was then performed on the area of interest with the laser at 50-100% power and an exposure time of 100 ms. After photobleaching, signal was measured with time-lapse image stacks obtained continuously using the 488 nm illumination laser.

### Determination of spindle length for pre-anaphase vs anaphase B

Ten monopolar spindles each for *klp5*+ and *klp5*Δ cells were identified which underwent rapid elongation after a period of a constant, stable length. The average length at this transition for *klp5*+ spindles was 2.40 *μ*m, and for *klp5*Δ spindles was 5.16 *μ*m.

### Dataset organization and kymograph creation

For all bipolar tracked data in this study, strains with Cut7-GFP or Cut7-3GFP, and *cut11*+ or *cut11-7* were pooled to create the kymographs and associated datasets displayed in figures 2-5. In addition, PA-GFP and fluorescence photobleaching experiments were pooled with all non-PA-GFP and unbleached experiments to create datasets for figures 2-3. The kymographs used for the quantification in figure 2 were compiled from strains MB951, MB1030, MB1032, MB1062, MB1064, MB1077, and MB1085 (table S4). The kymographs quantified for figure 3 were compiled from strains MB951 and MB1088 (table 4). The monopolar spindles analyzed in figure 4 graphs were compiled from strain MB1134. The bipolar spindles analyzed in figure 5 graphs were compiled from strains MB1131 and MB1155 and compared with the compiled data from figure 2.

All confocal images were transferred as Nikon nd2 files into the Fiji version of ImageJ (National Institutes of Health, Bethesda, MD) and displayed as pixel-interpolated, maximum-intensity projections. The FIJI plugin StackReg was utilized to align the data using the Transformation-Rigid Body option. The Image J segmented line tool was used to draw a straight line through the aligned spindle data. Finally, the Image J Reslice function with an output spacing of 1 *μ*m was used to create the kymographs shown.

### Track and SPB annotation

Annotations were made directly on the kymograph. We created in-house MATLAB software that allowed a user to interactively draw Cut7p tracks (Fig. S2). Some of these tracks were identified as corresponding to spindle-pole-localized Cut7-GFP, while other tracks were used for motility analysis. For bipolar spindle kymographs, both spindle poles were identified by areas of highest brightness throughout the kymograph. Two SPB lines were traced over those areas. Clusters of Cut7-GFP moving along spindle microtubules were then traced. The nearest SPB was used as a reference for describing poleward or anti-poleward movement. For monopolar spindle kymographs, the SPB was identified by the area of highest brightness throughout the kymograph, and an SPB line was traced. Clusters of Cut7-GFP moving poleward or antipoleward were then traced.

### Track merging and event identification

We started by merging tracks that were deemed to belong to the same global track. For example, if track 2 began within some time interval of track 1 ending, and the start position of track 2 was close to the end position of track 1, then we merged the tracks (Fig. S3). After a phase of merging, we identified a number of tracks with both poleward and anti-poleward motion. The tracks were then analyzed by splitting them into poleward, anti-poleward, and paused segments or events. Event identification was based on the average local velocity along the track. A cutoff value of 3 nm/sec was used to distinguish directed movement from pausing. Points with velocity less than the negative of the cutoff are assigned as poleward, while points with velocity greater than the cutoff are assigned as antipoleward. The remaining points were assigned as paused.

### Track analysis quantification

Tracks were analyzed with reference to the position of the SPB. To do that, we assigned each track to its nearest SPB. For monopolar spindles, there was only one pole and assignment was straightforward. For bipolar spindles with two SPBs, we picked the SPB that was closest to the track start position. Distance and velocity were measured relative to the closest SPB. For each kymograph, the frequency of directional events was calculated by counting the number of poleward and anti-poleward tracks and dividing by the total time of the kymograph. The switching frequency was calculated by counting the number of switches out of a state (poleward and anti-poleward movement) divided by the total time of all tracks in that state. For example, the switching frequency for exiting the poleward-moving state was calculated by dividing the number of transitions out of the poleward state by the total time of all poleward tracks.

## Acknowledgements

We thank Mohan Balasubramanian, Anne Paoletti, Takashi Toda, and Yoshi Wantanabe for gifts of strains used to construct the strains in this study, the National Bio-Resource Project – Yeast, Japan for providing NBRP strains FY23688 and FY31411, along with Pombase and the Wellcome Trust for providing online resources for Cut7p and other fission yeast proteins. We acknowledge the Department of Moleculer, Cellular, and Developmental Biology (MCDB) Light Microscopy Core Facility for assistance with live-cell imaging and the Biocore Facility of MCD for use of the Nanodrop 2000.

## Competing interests

The authors report no completing interests.

## Contribution

Conceptualization, MDB; Methodology, ZRG, MHJ, JRM, MDB; Software, SA; Validation, ZRG, SA, MHJ; Formal analysis, ZRG, SA, MDB; Investigation, ZRG, SA, MHJ; Data curation, ZRG, SA, MHJ; Writing - original draft preparation, ZRG, JRM, MDB; Writing - reviewing and editing, ZRG, SA, MHJ, JRM, MDB; Visualization, ZG, SA, MJ, MDB; Supervision, MDB; Project administration, MDB; Funding acquisition, MDB

## Funding

This work was supported by NIH grant R01 GM124371 and NSF grant DMR 1725065 to M.D.B.

## Data availability

All data and computer code used in this study are available upon request.

## Supplementary

See supplementary figures and tables in the attched file.

## Notes

### Competing Interest Statement

The authors have declared no competing interest.

## REFERENCES

Seyda Acar, David B. Carlson, Madhu S. Budamagunta, Vladimir Yarov-Yarovoy, John J. Correia, Milady R. Niñonuevo, Weitao Jia, Li Tao, Julie A. Leary, John C. Voss, James E. Evans, and Jonathan M. Scholey. The bipolar assembly domain of the mitotic motor kinesin-5. Nature Communications, 4:ncomms2348, January 2013.

Takashi Akera, Yuhei Goto, Masamitsu Sato, Masayuki Yamamoto, and Yoshinori Watanabe. Mad1 promotes chromosome congression by anchoring a kinesin motor to the kinetochore. Nature Cell Biology, 17(9):1124–1133, September 2015.

Rachel Avunie-Masala, Natalia Movshovich, Yael Nissenkorn, Adina Gerson-Gurwitz, Vladimir Fridman, Mardo Kõivomägi, Mart Loog, M. Andrew Hoyt, Arieh Zaritsky, and Larisa Gheber. Phospho-regulation of kinesin-5 during anaphase spindle elongation. Journal of Cell Science, 124:873–878, March 2011.

Sai K. Balchand, Barbara J. Mann, Janel Titus, Jennifer L. Ross, and Patricia Wadsworth. TPX2 Inhibits Eg5 by Interactions with Both Motor and Microtubule. Journal of Biological Chemistry, 290(28):17367–17379, July 2015.

Robert Blackwell, Oliver Sweezy-Schindler, Christopher Baldwin, Loren E. Hough, Matthew A. Glaser, and M. D. Betterton. Microscopic origins of anisotropic active stress in motor-driven nematic liquid crystals. Soft Matter, 12:2676–2687, January 2016.

Robert Blackwell, Oliver Sweezy-Schindler, Christopher Edelmaier, Zachary R. Gergely, Patrick J. Flynn, Salvador Montes, Ammon Crapo, Alireza Doostan, J. Richard McIntosh, Matthew A. Glaser, and Meredith D. Betterton. Contributions of Microtubule Dynamic Instability and Rotational Diffusion to Kinetochore Capture. Biophysical Journal, 112(3):552–563, February 2017.

Anne Blangy, Heidi A. Lane, Pierre d’Hérin, Maryannick Harper, Michel Kress, and Erich A. Nigg. Phosphorylation by p34cdc2 regulates spindle association of human Eg5, a kinesin-related motor essential for bipolar spindle formation in vivo. Cell, 83(7):1159–1169, December 1995.

Tatyana Bodrug, Elizabeth M Wilson-Kubalek, Stanley Nithianantham, Alex F Thompson, April Alfieri, Ignas Gaska, Jennifer Major, Garrett Debs, Sayaka Inagaki, Pedro Gutierrez, Larisa Gheber, Richard J McKenney, Charles Vaughn Sindelar, Ronald Milligan, Jason Stumpff, Steven S Rosenfeld, Scott T Forth, and Jawdat Al-Bassam. The kinesin-5 tail domain directly modulates the mechanochemical cycle of the motor domain for anti-parallel microtubule sliding. eLife, 9:e51131, January 2020.

Mishan Britto, Adeline Goulet, Syeda Rizvi, Ottilie von Loeffelholz, Carolyn A. Moores, and Robert A. Cross. Schizosaccharomyces pombe kinesin-5 switches direction using a steric blocking mechanism. Proceedings of the National Academy of Sciences, page 201611581, November 2016.

J. Cahu and T. Surrey. Motile microtubule crosslinkers require distinct dynamic properties for correct functioning during spindle organization in Xenopus egg extract. Journal of Cell Science, 122(9):1295–1300, April 2009.

Julie Cahu, Aurelien Olichon, Christian Hentrich, Henry Schek, Jovana Drinjakovic, Cunjie Zhang, Amanda Doherty-Kirby, Gilles Lajoie, and Thomas Surrey. Phosphorylation by Cdk1 Increases the Binding of Eg5 to Microtubules In Vitro and in Xenopus Egg Extract Spindles. PLOS ONE, 3(12):e3936, December 2008.

D. G. Cole, W. M. Saxton, K. B. Sheehan, and J. M. Scholey. A “slow” homotetrameric kinesin-related motor protein purified from Drosophila embryos. Journal of Biological Chemistry, 269(37):22913–22916, September 1994.

R. Ding, K. L. McDonald, and J. R. McIntosh. Three-dimensional reconstruction and analysis of mitotic spindles from the yeast, Schizosaccharomyces pombe. The Journal of Cell Biology, 120(1):141–151, January 1993.

D. R. Drummond and I. M. Hagan. Mutations in the bimC box of Cut7 indicate divergence of regulation within the bimC family of kinesin related proteins. Journal of Cell Science, 111(7):853–865, April 1998.

André Düselder, Vladimir Fridman, Christina Thiede, Alice Wiesbaum, Alina Goldstein, Dieter R. Klopfenstein, Olga Zaitseva, Marcel E. Janson, Larisa Gheber, and Christoph F. Schmidt. Deletion of the Tail Domain of the Kinesin-5 Cin8 Affects Its Directionality. Journal of Biological Chemistry, 290(27):16841–16850, July 2015.

M. Edamatsu. Molecular properties of the N-terminal extension of the fission yeast kinesin-5, Cut7. Genetics and molecular research : GMR, 15(1), February 2016.

Masaki Edamatsu. Bidirectional motility of the fission yeast kinesin-5, Cut7. Biochemical and Biophysical Research Communications, 446(1):231–234, March 2014.

Christopher Edelmaier, Adam R Lamson, Zachary R Gergely, Saad Ansari, Robert Blackwell, J Richard McIntosh, Matthew A Glaser, and Meredith D Betterton. Mechanisms of chromosome biorientation and bipolar spindle assembly analyzed by computational modeling. eLife, 9:e48787, February 2020.

A. P. Enos and N. R. Morris. Mutation of a gene that encodes a kinesin-like protein blocks nuclear division in A. nidulans. Cell, 60(6):1019–1027, March 1990.

Vladimir Fridman, Adina Gerson-Gurwitz, Ofer Shapira, Natalia Movshovich, Stefan Lakämper, Christoph F. Schmidt, and Larisa Gheber. Kinesin-5 Kip1 is a bi-directional motor that stabilizes microtubules and tracks their plus-ends in vivo. J Cell Sci, 126(18):4147–4159, September 2013.

Chuanhai Fu, Jonathan J. Ward, Isabelle Loiodice, Guilhem Velve-Casquillas, Francois J. Nedelec, and Phong T. Tran. Phospho-Regulated Interaction between Kinesin-6 Klp9p and Microtubule Bundler Ase1p Promotes Spindle Elongation. Developmental Cell, 17(2):257–267, August 2009.

Alyssa Gable, Minhua Qiu, Janel Titus, Sai Balchand, Nick P. Ferenz, Nan Ma, Elizabeth S. Collins, Carey Fagerstrom, Jennifer L. Ross, Ge Yang, and Patricia Wadsworth. Dynamic reorganization of Eg5 in the mammalian spindle throughout mitosis requires dynein and TPX2. Molecular Biology of the Cell, 23(7): 1254–1266, April 2012.

Zachary R. Gergely, Ammon Crapo, Loren E. Hough, J. Richard McIntosh, and Meredith D. Betterton. Kinesin-8 effects on mitotic microtubule dynamics contribute to spindle function in fission yeast. Molecular Biology of the Cell, 27(22):3490–3514, November 2016.

Adina Gerson-Gurwitz, Christina Thiede, Natalia Movshovich, Vladimir Fridman, Maria Podolskaya, Tsafi Danieli, Stefan Lakämper, Dieter R. Klopfenstein, Christoph F. Schmidt, and Larisa Gheber. Directionality of individual kinesin-5 Cin8 motors is modulated by loop 8, ionic strength and microtubule geometry. The EMBO Journal, 30(24):4942, December 2011.

Donna M. Gordon and David M. Roof. The Kinesin-related Protein Kip1p of Saccharomyces cerevisiae Is Bipolar. Journal of Biological Chemistry, 274(40): 28779–28786, October 1999.

Gohta Goshima and Jonathan M. Scholey. Control of Mitotic Spindle Length. Annual Review of Cell and Developmental Biology, 26(1):21–57, 2010.

Iain Hagan and Mitsuhiro Yanagida. Novel potential mitotic motor protein encoded by the fission yeast cut7+ gene. Nature, 347(6293):563–566, October 1990.

Iain Hagan and Mitsuhiro Yanagida. Kinesin-related cut 7 protein associates with mitotic and meiotic spindles in fission yeast. Nature, 356(6364):74, March 1992.

M. M. Heck, A. Pereira, P. Pesavento, Y. Yannoni, A. C. Spradling, and L. S. Goldstein. The kinesin-like protein KLP61F is essential for mitosis in Drosophila. The Journal of Cell Biology, 123(3):665–679, November 1993.

Emily R. Hildebrandt, Larisa Gheber, Tami Kingsbury, and M. Andrew Hoyt. Homotetrameric Form of Cin8p, a Saccharomyces cerevisiae Kinesin-5 Motor, Is Essential for Its in Vivo Function. Journal of Biological Chemistry, 281(36):26004–26013, September 2006.

Tatsuya Hirano, Shin-ichi Funahashi, Tadashi Uemura, and Mitsuhiro Yanagida. Isolation and characterization of Schizosaccharomyces pombe cutmutants that block nuclear division but not cytokinesis. The EMBO Journal, 5(11):2973–2979, November 1986.

M. A. Hoyt, L. He, K. K. Loo, and W. S. Saunders. Two Saccharomyces cerevisiae kinesin-related gene products required for mitotic spindle assembly. The Journal of Cell Biology, 118(1):109–120, July 1992.

Woosang Hwang, Takashi Toda, and Masashi Yukawa. Complementation of fission yeast kinesin-5/Cut7 with human Eg5 provides a versatile platform for screening of anticancer compounds. Bioscience, Biotechnology, and Biochemistry, 86(2):254–259, February 2022.

Iana Kalinina, Amitabha Nandi, Petrina Delivani, Mariola R. Chacón, Anna H. Klemm, Damien Ramunno-Johnson, Alexander Krull, Benjamin Lindner, Nenad Pavin, and Iva M. Tolić-Norrelykke. Pivoting of microtubules around the spindle pole accelerates kinetochore capture. Nature Cell Biology, 2012.

Lukas C. Kapitein, Erwin J. G. Peterman, Benjamin H. Kwok, Jeffrey H. Kim, Tarun M. Kapoor, and Christoph F. Schmidt. The bipolar mitotic kinesin Eg5 moves on both microtubules that it crosslinks. Nature, 435(7038):114–118, May 2005.

A. S. Kashina, R. J. Baskin, D. G. Cole, K. P. Wedaman, W. M. Saxton, and J. M. Scholey. A bipolar kinesin. Nature, 379(6562):270–272, January 1996.

Lara Katharina Krüger, Matthieu Gélin, Liang Ji, Carlos Kikuti, Anne Houdusse, Manuel Théry, Laurent Blanchoin, and Phong T Tran. Kinesin-6 Klp9 orchestrates spindle elongation by regulating microtubule sliding and growth. eLife, 10:e67489, June 2021.

Adam R. Lamson, Jeffrey M. Moore, Fang Fang, Matthew A. Glaser, Michael Shelley, and Meredith D. Betterton. Comparison of explicit and mean-field models of cytoskeletal filaments with crosslinking motors. arXiv:2011.08156 [physics], November 2020.

Andrei Lupas, Marc Van Dyke, and Jeff Stock. Predicting Coiled Coils from Protein Sequences. Science, 252(5009):1162–1164, May 1991.

Nan Ma, U. S. Tulu, Nick P. Ferenz, Carey Fagerstrom, Andrew Wilde, and Patricia Wadsworth. Poleward Transport of TPX2 in the Mammalian Mitotic Spindle Requires Dynein, Eg5, and Microtubule Flux. Molecular Biology of the Cell, 21(6):979–988, March 2010.

Nan Ma, Janel Titus, Alyssa Gable, Jennifer L. Ross, and Patricia Wadsworth. TPX2 regulates the localization and activity of Eg5 in the mammalian mitotic spindle. J Cell Biol, 195(1):87–98, October 2011.

Barbara J. Mann and Patricia Wadsworth. Kinesin-5 Regulation and Function in Mitosis. Trends in Cell Biology, September 2018.

H. Masuda, M. Sevik, and W. Z. Cande. In vitro microtubule-nucleating activity of spindle pole bodies in fission yeast Schizosaccharomyces pombe: Cell cycle-dependent activation in xenopus cell-free extracts. The Journal of Cell Biology, 117(5):1055–1066, June 1992.

Thomas U. Mayer, Tarun M. Kapoor, Stephen J. Haggarty, Randall W. King, Stuart L. Schreiber, and Timothy J. Mitchison. Small Molecule Inhibitor of Mitotic Spindle Bipolarity Identified in a Phenotype-Based Screen. Science, 286(5441):971–974, October 1999.

J. Richard McIntosh and Story C. Landis. THE DISTRIBUTION OF SPINDLE MICROTUBULES DURING MITOSIS IN CULTURED HUMAN CELLS. Journal of Cell Biology, 49(2):468–497, May 1971.

Sergio Moreno, Amar Klar, and Paul Nurse. Molecular genetic analysis of fission yeast Schizosaccharomyces pombe. In Guide to Yeast Genetics and Molecular Biology, volume Volume 194, pages 795–823. 1991.

Makoto J. Ohira, David G. Hendrickson, R. Scott McIsaac, and Nicholas Rhind. An Estradiol-Inducible Promoter Enables Fast, Graduated Control of Gene Expression in Fission Yeast. Yeast, pages n/a–n/a, January 2017.

Koei Okazaki, Noriko Okazaki, Kazuhiko Kume, Shigeki Jinno, Koichi Tanaka, and Hiroto Okayama. High-frequency transformation method and library transducing vectors for cloning mammalian cDNAs by trans-complementation of Schizosaccharomyces pombe. Nucleic Acids Research, 18(22):6485–6489, November 1990.

Zachary T. Olmsted, Andrew G. Colliver, Timothy D. Riehlman, and Janet L. Paluh. Kinesin-14 and kinesin-5 antagonistically regulate microtubule nucleation by γ-TuRC in yeast and human cells. Nature Communications, 5:5339, October 2014.

A L Pidoux, M LeDizet, and W Z Cande. Fission yeast pkl1 is a kinesin-related protein involved in mitotic spindle function. Molecular Biology of the Cell, 7(10): 1639–1655, October 1996.

Joseph Rapley, Marta Nicolàs, Aaron Groen, Laura Regué, M. Teresa Bertran, Carme Caelles, Joseph Avruch, and Joan Roig. The NIMA-family kinase Nek6 phosphorylates the kinesin Eg5 at a novel site necessary for mitotic spindle formation. Journal of Cell Science, 121(23):3912–3921, December 2008.

Sergio A. Rincon, Adam Lamson, Robert Blackwell, Viktoriya Syrovatkina, Vincent Fraisier, Anne Paoletti, Meredith D. Betterton, and Phong T. Tran. Kinesin-5-independent mitotic spindle assembly requires the antiparallel microtubule crosslinker Ase1 in fission yeast. Nature Communications, 8:15286, May 2017.

Adrianna S. Rodriguez, Alison N. Killilea, Joseph Batac, Jason Filopei, Dimitre Simeonov, Ida Lin, and Janet L. Paluh. Protein complexes at the microtubule organizing center regulate bipolar spindle assembly. Cell Cycle, 7(9):1246–1253, May 2008.

D M Roof, P B Meluh, and M D Rose. Kinesin-related proteins required for assembly of the mitotic spindle. Journal of Cell Biology, 118(1):95–108, July 1992.

Johanna Roostalu, Christian Hentrich, Peter Bieling, Ivo A. Telley, Elmar Schiebel, and Thomas Surrey. Directional Switching of the Kinesin Cin8 Through Motor Coupling. Science, 332(6025):94–99, April 2011.

K. E. Sawin and T. J. Mitchison. Mutations in the kinesin-like protein Eg5 disrupting localization to the mitotic spindle. Proceedings of the National Academy of Sciences, 92(10):4289–4293, May 1995.

Kenneth E. Sawin, Katherine LeGuellec, Michel Philippe, and Timothy J. Mitchison. Mitotic spindle organization by a plus-end-directed microtubule motor. Nature, 359(6395):540–543, October 1992.

Jessica E. Scholey, Stanley Nithianantham, Jonathan M. Scholey, and Jawdat Al-Bassam. Structural basis for the assembly of the mitotic motor Kinesin-5 into bipolar tetramers. eLife, 3:e02217, April 2014.

Jonathan M. Scholey, Gul Civelekoglu-Scholey, and Ingrid Brust-Mascher. Anaphase B. Biology, 5(4):51, December 2016.

Ofer Shapira and Larisa Gheber. Motile properties of the bi-directional kinesin-5 Cin8 are affected by phosphorylation in its motor domain. Scientific Reports, 6: 25597, May 2016.

Ofer Shapira, Alina Goldstein, Jawdat Al-Bassam, and Larisa Gheber. Possible physiological role for bi-directional motility and motor clustering of the mitotic kinesin-5 Cin8. J Cell Sci, page jcs.195040, January 2017.

David J. Sharp, Kent L. McDonald, Heather M. Brown, Heinrich J. Matthies, Claire Walczak, Ron D. Vale, Timothy J. Mitchison, and Jonathan M. Scholey. The Bipolar Kinesin, KLP61F, Cross-links Microtubules within Interpolar Microtubule Bundles of Drosophila Embryonic Mitotic Spindles. The Journal of Cell Biology, 144(1):125–138, January 1999.

Yuta Shimamoto, Scott Forth, and Tarun M. Kapoor. Measuring Pushing and Braking Forces Generated by Ensembles of Kinesin-5 Crosslinking Two Microtubules. Developmental Cell, 34(6):669–681, September 2015.

Sudhir Kumar Singh, Himanshu Pandey, Jawdat Al-Bassam, and Larisa Gheber. Bidirectional motility of kinesin-5 motor proteins: Structural determinants, cumulative functions and physiological roles. Cellular and Molecular Life Sciences, 75(10):1757–1771, May 2018.

Hilary A. Snaith, Andreas Anders, Itaru Samejima, and Kenneth E. Sawin. Chapter 9 - New and Old Reagents for Fluorescent Protein Tagging of Microtubules in Fission Yeast: Experimental and Critical Evaluation. In Lynne Cassimeris and Phong Tran, editor, Methods in Cell Biology, volume Volume 97, pages 147–172. 2010.

Li Tao, Alex Mogilner, Gul Civelekoglu-Scholey, Roy Wollman, James Evans, Henning Stahlberg, and Jonathan M. Scholey. A Homotetrameric Kinesin-5, KLP61F, Bundles Microtubules and Antagonizes Ncd in Motility Assays. Current Biology, 16(23):2293–2302, December 2006.

Christina Thiede, Vladimir Fridman, Adina Gerson-Gurwitz, Larisa Gheber, and Christoph F. Schmidt. Regulation of bi-directional movement of single kinesin-5 Cin8 molecules. BioArchitecture, 2(2):70–74, March 2012.

Cynthia L. Troxell, Mark A. Sweezy, Robert R. West, Karen D. Reed, Bryan D. Carson, Alison L. Pidoux, W. Zacheus Cande, and J. Richard McIntosh. Pkl1 +andklp2 +: Two Kinesins of the Kar3 Subfamily in Fission Yeast Perform Different Functions in Both Mitosis and Meiosis. Molecular Biology of the Cell, 12(11):3476–3488, November 2001.

A. Unsworth, H. Masuda, S. Dhut, and T. Toda. Fission yeast kinesin-8 Klp5 and Klp6 are interdependent for mitotic nuclear retention and required for proper microtubule dynamics. Molecular Biology of the Cell, 19(12):5104–5115, 2008.

Marianne Uteng, Christian Hentrich, Kota Miura, Peter Bieling, and Thomas Surrey. Poleward transport of Eg5 by dynein–dynactin in Xenopus laevis egg extract spindles. The Journal of Cell Biology, 182(4):715–726, August 2008.

Siet M. J. L. van den Wildenberg, Li Tao, Lukas C. Kapitein, Christoph F. Schmidt, Jonathan M. Scholey, and Erwin J. G. Peterman. The Homotetrameric Kinesin-5 KLP61F Preferentially Crosslinks Microtubules into Antiparallel Orientations. Current Biology, 18(23):1860–1864, December 2008.

Jonathan J. Ward, Hélio Roque, Claude Antony, and François Nédélec. Mechanical design principles of a mitotic spindle. eLife, 3:e03398, January 2015.

J. S. Weinger, M. Qiu, G. Yang, and T. M. Kapoor. A nonmotor microtubule binding site in kinesin-5 is required for filament crosslinking and sliding. Current Biology, 21(2):154–160, 2011.

R. R. West, T. Malmstrom, C. L. Troxell, and J. R. McIntosh. Two Related Kinesins, klp5+ and klp6+, Foster Microtubule Disassembly and Are Required for Meiosis in Fission Yeast. Molecular Biology of the Cell, 12(12):3919–3932, 2001.

Robert R. West, Elena V. Vaisberg, Rubai Ding, Paul Nurse, and J. Richard McIntosh. Cut11 +: A Gene Required for Cell Cycle-dependent Spindle Pole Body Anchoring in the Nuclear Envelope and Bipolar Spindle Formation in Schizosaccharomyces pombe. Molecular Biology of the Cell, 9(10):2839–2855, October 1998.

Y. Yamagishi, C. H. Yang, Y. Tanno, and Y. Watanabe. MPS1/Mph1 phosphorylates the kinetochore protein KNL1/Spc7 to recruit SAC components. Nature Cell Biology, 14(7):746–752, 2012.

Masashi Yukawa, Chiho Ikebe, and Takashi Toda. The Msd1–Wdr8–Pkl1 complex anchors microtubule minus ends to fission yeast spindle pole bodies. J Cell Biol, 209(4):549–562, May 2015.

Masashi Yukawa, Yusuke Yamada, Tomoaki Yamauchi, and Takashi Toda. Two spatially distinct Kinesin-14 Pkl1 and Klp2 generate collaborative inward forces against Kinesin-5 Cut7 in S. pombe. J Cell Sci, page jcs.210740, 2018.

Masashi Yukawa, Masaki Okazaki, Yasuhiro Teratani, Ken’ya Furuta, and Takashi Toda. Kinesin-6 Klp9 plays motor-dependent and -independent roles in collaboration with Kinesin-5 Cut7 and the microtubule crosslinker Ase1 in fission yeast. Scientific Reports, 9(1):7336, May 2019.

Masashi Yukawa, Yasuhiro Teratani, and Takashi Toda. How Essential Kinesin-5 Becomes Non-Essential in Fission Yeast: Force Balance and Microtubule Dynamics Matter. Cells, 9(5):1154, May 2020.

Dan Zhang and Snezhana Oliferenko. Tts1, the fission yeast homologue of the TMEM33 family, functions in NE remodeling during mitosis. Molecular Biology of the Cell, 25(19):2970–2983, October 2014.

